# Oncohistone inhibition reshapes tumor–microenvironment communication in Diffuse Midline Glioma (DMG)

**DOI:** 10.64898/2026.06.17.731637

**Authors:** Niloofar Khairkhah, Mostafa M. H. Ibrahim, Sienna L. Galban, Megan Faunce, Lily Rober, Carlen baker, Robert Doherty, Rodrigo Cartaxo, Carl Koschmann, Yue Zhao, Stefanie Galban

**Affiliations:** Department of Radiology, The University of Michigan Medical School, Ann Arbor, MI 48109, United States; Davidson School of Chemical Engineering, Purdue University, West Lafayette, IN, United States; Department of Pediatrics, University of Michigan Medical School, Ann Arbor, Michigan, USA; Department of Neurology, University of Michigan Medical School, Ann Arbor, Michigan, USA; Gilbert S. Omenn Department of Computational Medicine and Bioinformatics, The University of Michigan Medical School, Ann Arbor, MI 48109, United States; Department of Surgery, The University of Michigan Medical School, Ann Arbor, MI 48109, United States; Center for Molecular Imaging, The University of Michigan Medical School, Ann Arbor, MI 48109, United States; Rogel Cancer Center, The University of Michigan Medical School, Ann Arbor, MI 48109, United States

**Keywords:** DMG, oncohistone inhibition, Tumor microenvironment, neuron-glioma interaction, neuron-neuron interaction

## Abstract

**Background:** Diffuse midline glioma (DMG) is a lethal pediatric brain tumor driven by the H3K27M oncohistone, which disrupts epigenetic regulation and promotes tumor proliferation. While prior studies show that H3K27M is essential for tumor initiation, its role in established tumors, tumor microenvironment (TME) regulation, and therapeutic response remain unclear.

**Methods:** Here, we developed inducible and reversible H3.3K27M and H3.1K27M cell and mouse models to study oncohistone-dependent effects and the immune/stromal microenvironment. We generated a tetracycline-inducible PiggyBac-based oncohistone expression cassette in patient- and murine-derived models and validated inducible and reversible H3K27M expression.

**Results:** Re-expression of H3K27M in knockout cells induced morphological changes and suppressed astrocytic markers. Chromatin accessibility profiling revealed locus-specific differences between ON, OFF, and OFF–ON, including changes at loci associated with immune regulation and tumor–microenvironment interactions. Single-cell RNA sequencing demonstrated that the oncohistone reshapes the TME. H3K27M expression promotes tumor–neuron interactions, enhances neuronal excitability, excitatory/inhibitory imbalance, and synaptic connectivity that likely supports tumor proliferation. These effects are associated with increased glutamatergic signaling and enhanced tumor–neuron coupling through glutamate transport and receptor pathways, including EAAT1 (*SLC1A3*) and AMPARs (*GRIA3*). Conversely, H3K27M inhibition reduces neuronal excitation, disrupts tumor-associated signaling, and partially restores neuron–neuron and neuron–immune communications. These findings identify H3K27M as a key driver of excitatory neuron-to-tumor coupling and immunosuppression in DMG.

**Conclusions:** Overall, our findings demonstrate that H3K27M extensively reshapes TME in DMG and support direct oncohistone targeting as a potential therapeutic strategy, including potential CRISPR-based or small-molecule approaches for patients with H3K27M-mutant DMG.

**Key Points:**

- We developed inducible and reversible H3K27M DMG models to investigate the role of H3K27M in the tumor microenvironment.
- H3K27M promotes tumor–neuron communication, while its inhibition disrupts these interactions, supporting H3K27M-targeted therapies for DMG.

**Importance of Study:** Diffuse midline glioma (DMG) remains one of the deadliest pediatric brain tumors, with limited effective treatment options and poor patient survival. Although the H3K27M oncohistone is recognized as a key driver of tumor initiation, its role in maintaining tumor progression and shaping the tumor microenvironment is unclear. In this study, we developed inducible and reversible H3.3K27M and H3.1K27M murine and patient-derived DMG cell- and mouse-models that enabled precise control of the oncohistone expression. Using these models, we demonstrate that H3K27M actively promotes tumor–neuron interactions, neuronal excitability, and glutamatergic signaling pathways that support tumor growth. Importantly, inhibition of H3K27M disrupted these tumor-associated signaling networks and partially restored neuron– immune communication within the tumor microenvironment. Together, these findings demonstrate that H3K27M extensively reshapes the tumor microenvironment in these Diffuse Midline Gliomas and provides strong rationale for directly targeting the oncohistone as a therapeutic strategy for patients with H3K27M-mutant DMG.

## Introduction

Diffuse Midline Glioma (DMG) remain one of the most devastating and challenging to treat pediatric brain cancers ^1^. These tumors arise in the brainstem, are highly infiltrative which makes surgical resection impossible ^2^. Radiotherapy remains the standard of care, but it provides only temporary symptom relief ^3,4^. Despite extensive clinical efforts, most patients experience rapid tumor progression, and treatment options remain limited, especially at recurrence, where agents such as ONC201 (dordaviprone or Modeyso™), have been explored ^5,6^. The consistent failure of existing therapies underscores a fundamental gap in our understanding of the pathobiology and therapeutic resistance mechanisms that drive this intractable disease. This highlights a critical need to better understand the biology of DMG and identify more effective treatment strategies.

The hallmark genetic alteration in DMG is the H3K27M mutation, which disrupts epigenetic regulation by inhibiting PRC2 activity, resulting in global loss of H3K27me3, increased H3K27 acetylation, and maintenance of stem-like tumor states ^7,8^. Given the central role of H3.3K27M in tumor maintenance, efforts to therapeutically target this mutation are ongoing, including strategies such as antisense oligonucleotides (ASOs) ^9^. Several studies have investigated the role of H3K27M by genetic deletion or silencing of the mutant allele in patient-derived models, including Clustered Regularly Interspaced Short Palindromic Repeats (CRISPR)/CRISPR-associated protein 9 (Cas9)-knockout and shRNA-based knockdown approaches ^8,10^. These studies demonstrated that DMG cells are dependent on H3K27M, as loss of the mutant allele abolishes tumorigenic potential *in vivo* ^8^. Building on these models, we previously used isogenic H3.3K27M and H3.3K27M-KO cell pairs to demonstrate that H3K27M drives widespread transcriptional changes in DMG cells ^11^. These findings suggest that targeting H3.3K27M could represent a promising therapeutic strategy alone and in combination with standard of care (IR) ^11^. However, studies which uses isogenic pairs primarily address tumor initiation and do not reflect the clinical setting, where tumors are already established at diagnosis. Emerging evidence further indicates that the neuronal microenvironment plays a key role in DMG progression ^12^. DMG arises from oligodendroglial lineage precursor cells (OPCs) ^13,14^, which in the normal brain, during postnatal development and adulthood, actively communicate with neurons through both paracrine signaling and direct synaptic interactions, including glutamatergic and GABAergic neuron-to-OPC synapses ^15^. This highlights a close functional relationship between tumor-initiating and the neuronal niche. However, it remains unclear whether H3.3K27M inhibition in established tumors can induce tumor regression, alter tumor–microenvironment (TME) interactions, or disrupt tumor–neuron communication. Given the limited success of therapies targeting downstream epigenetic and transcriptional programs, understanding the consequences of direct H3K27M inhibition is critical for future therapeutic development. Similar to genetic inhibition of oncogenic drivers such as oncogenic KRAS in pancreatic and lung cancers ^16,17^, this approach may help define the therapeutic potential of directly targeting H3K27M.

Here, we developed inducible and reversible H3.3K27M and H3.1K27M cell and mouse models for DMG that allow precise control of oncohistone expression in cells and *in vivo*. This allowed us to assess the effects of oncohistone activation and inactivation on tumor growth and define the associated cellular and epigenetic changes and their impact on the tumor microenvironment (TME).

## Materials and Methods

### Cell culture and doxycycline-inducible H3.3K27M vector design

The human DMG isogenic pair including human SU-DIPG XIII parental (H3.3K27M), CRISPR edited SU-DIPG XIII K27M-KO isogenic pair, and BT245 H3.3K27M parental and CRISPR edited BT245 K27M-KO, kindly provided by Dr. Jabado (Pediatrics, McGill University) ^8^, and the murine DMG cell line UC-BL6-B7(PPK) and UC-BL6-D3(PPW) cell lines, kindly provided by Dr. Timothy Pheonix (University of Cincinnati) ^18^ were used in this study. We designed a PiggyBac donor construct in which H3.3 K27M and mCherry are co-expressed via an internal ribosome entry site (IRES) under the control of a tetracycline-responsive element (TRE). Detailed vector design and construct information are provided in the Supplementary Methods.

### Generation of inducible and reversible murine DMG cells using Intra uterine electroporation (IUE) model

The transposase helper plasmid pCAG-PBase, together with the donor constructs PB-CAG-PdgfraD824V-Ires-EGFP (PDGFRA D842V) and PB-CAG-DNp53-Ires-luciferase (dominant-negative TP53, referred to herein as TP53), were described previously and were used here without further modification ^19^. We also used PB TetON-TRE H3.3K27M-mCherry CAG rtTA to generate the inducible H3.3K27M IUE model. A tumor bearing mouse was harvested and processed to establish the PPiK tumors (P: PDGFRA, P: p53, iK: inducible H3.3K27M) cells. Detailed information is provided in the Supplementary Methods.

### Generation of inducible and reversible human and murine DMG cells

The iH3.3K27M PiggyBac donor construct was first transfected into HEK293T and then DMG cells including SU-DIPG XIII K27M-KO (here DIPG XIII KO), BT245 K27M-KO (here BT245 KO) and PPW cells. Stable integration was achieved by co-transfection with the pCAG-PBase transposase plasmid in TSM complete medium supplemented with 2 μg/mL dox. mCherry-positive cells were sorted and here referred to as HEK293T+iH3.3K27M, DIPG XIII KO+iH3.3K27M, BT245 KO+iH3.3K27M, and PPW+iH3.3K27M. Detailed information is provided in the Supplementary Methods.

### Live and Intracellular flow cytometry

We performed live and intracellular flow cytometry on three inducible cells, HEK293T+iH3.3K27M, BT245 KO+iH3.3K27M and PPiK(iH3.3K27M) under ON, OFF, and OFF–ON dox. H3.3K27M expression was assessed using mutant-specific H3K27M antibodies or isotype controls. Samples were acquired on ZE5 Cell Analyzer flow cytometer and analyzed using FlowJo v10.10. Manufacturer details for antibodies are listed in Supplementary Table1 and detailed information is provided in the Supplementary Methods.

### Western Blot

Total protein and histone protein extracts were prepared and analyzed by Western blotting as previously described ^20^. Membranes were probed with antibodies against GFAP, EYA4, H3.3K27M, total H4, and β-actin, followed by HRP-conjugated secondary antibodies. Detailed extraction procedures, antibody information, and blotting conditions are provided in the Supplementary Methods and Supplementary Table1.

### Neurosphere assay

PPiK(iH3.3K27M) cells were cultured under ON and OFF groups for neurosphere formation assays. Neurosphere size and number were quantified using ImageJ from three independent experiments. Detailed culture conditions and analysis methods are provided in the Supplementary Methods.

### Cell proliferation assay

PPW+iH3.3K27M cells were irradiated or left untreated, with ONC201 (dordaviprone or Modeyso™), or a combination of both in ON and OFF groups. Cell viability was measured using the CellTiter-Glo assay, and IC50 values were calculated using GraphPad Prism. See Supplementary Methods for details.

### Bulk transposase-accessible chromatin with sequencing (ATAC-Seq)

ATAC-seq was performed on BT245 KO+iH3.3K27M cells cultured under ON, OFF, and OFF– ON. Sequencing and bioinformatic analyses were performed by Active Motif (Carlsbad, CA). Detailed experimental and bioinformatic methods are provided in the Supplementary Methods.

### Generation of inducible and reversible H3.3K27M *in vivo* models

All animal procedures were approved by the Institutional Animal Care and Use Committee (IACUC) of University of Michigan and Unit for Laboratory Animal Medicine (ULAM) under approved protocol PRO00012162 to Stefanie Galban. Flank xenograft studies were performed using BT245 KO+iH3.3K27M cells implanted subcutaneously into NU/NU mice maintained on dox-containing water. For orthotopic studies, PPW+iH3.3K27M cells were stereotactically implanted into the brains of C57BL/6 mice as previously described ^21^. Mice were maintained on dox prior to implantation and subsequently randomized to ON or OFF groups. Tumor growth and survival were monitored as indicated. Detailed implantation procedures, treatment schedules, and animal numbers are provided in the Supplementary Methods. In utero electroporation (IUE) was performed in CD1 embryos, using the same plasmid mixture and electroporation protocol described above. All moms received dox-containing drinking water (0.5 mg/mL, in sterile water). Pups remained with dox water and screened by BLI.

### FLEX single cell RNA-sequencing (scRNA-seq)

IUE derived tumors and adjacent microenvironmental tissue were collected and processed by the University of Michigan Advanced Genomics Core using the 10x Genomics FLEX single-cell RNA sequencing. Detailed experimental procedures, sequencing parameters, and bioinformatic analyses are provided in the Supplementary Methods. All analysis code and figure-generation scripts are publicly available on <u>GitHub</u> (https://github.com/yzhao80/iH3.3K27M-DMG-scRNA-seq).

## Results

### Inducible and Reversible Oncohistone Expression in DMG cells and tumors

To establish inducible and reversible control of the oncohistone (H3.3K27M) expression, we designed a dox-responsive PiggyBac (PB) system (Fig. 1A). First, we generated a murine DMG cell line using an IUE model (Fig. 1B) ^22^, in which four PiggyBac plasmids (PBase, PDGFRA D842V, DNTP53, and the inducible H3.3K27M construct) were electroporated into developing neural progenitors of mouse embryos at E14. Tumor formation and progression were monitored by BLI (Fig. 1C). This strategy resulted in the formation of DMG tumors in which tumors were collected and dissociated to establish a neurosphere cell line PPiK(iH3.3K27M) (Fig. 1D). Oncohistone expression was monitored by mCherry fluorescence.

**Figure 1.**
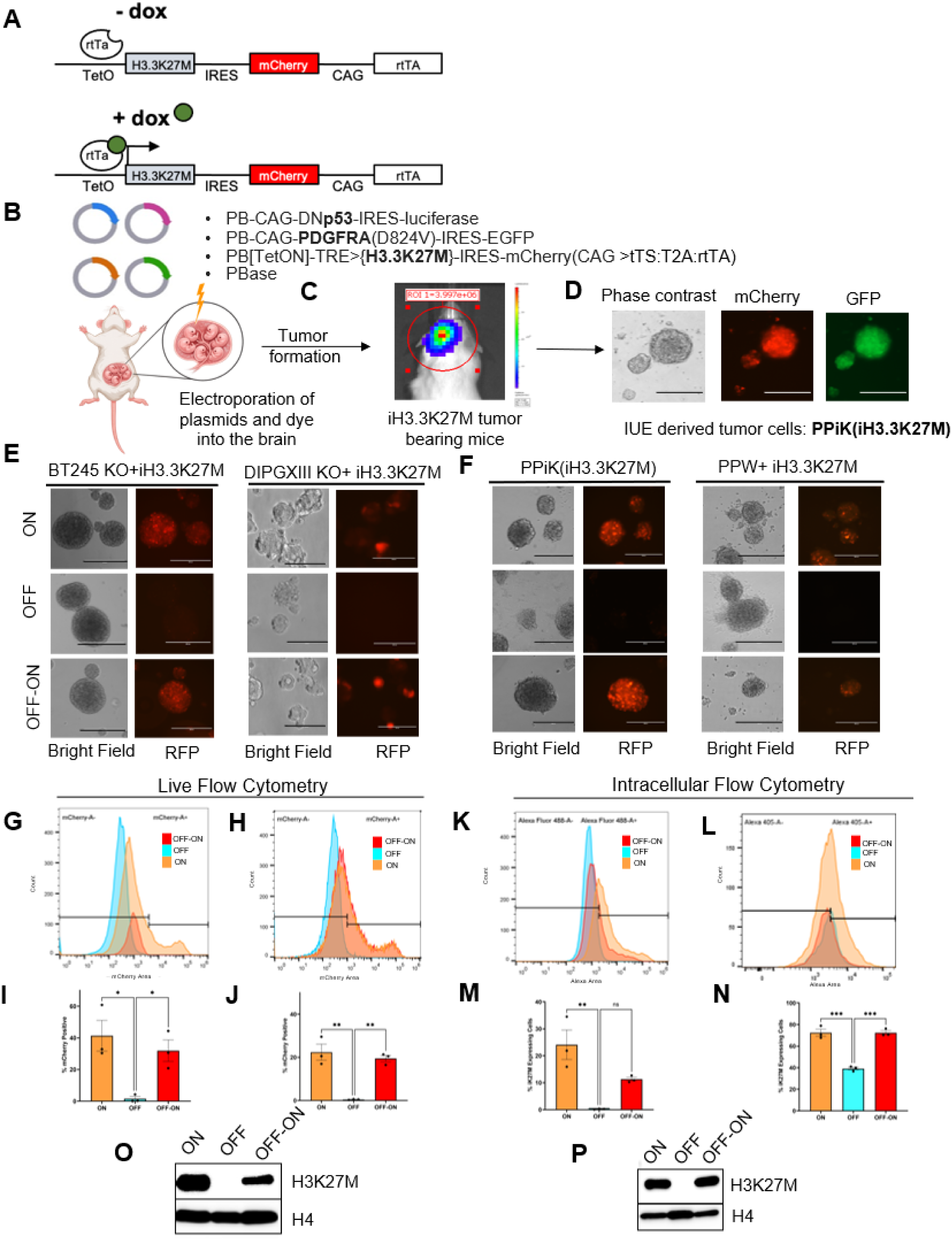
Generation of inducible and reversible H3.3K27M cell and mouse models. **A.** Schematic of the tetracycline-inducible construct: the ubiquitous CAG promoter drives expression of reverse tetracycline trans-activator (rtTA), which can bind to the TetO sites within the Tetracycline response element (TRE) upon dox induction to activate the expression of H3.3K27M and IRES dependent mCherry. **B.** PiggyBac vectors and intra uterine electroporation (IUE): at E13.5-14.5, embryos were removed from a pregnant CD1 mouse, a mix of four plasmids and tracer dye were injected into the fourth ventricle with a glass capillary prior to electroporation. **C.** Representative bioluminescence imaging (BLI) image depicting ability to monitor tumor development. **D.** Establishment of neutrosphere cell culture (PPiK(iH3.3K27M)) from tumors collected and dissociated approximately 30 days after birth and representative phase contrast and fluorescent images. **E. -F.** Representative phase contrast and fluorescence images of human (BT245 KO+iH3.3K27M, DIPGXIII KO+iH3.3K27M), and murine DMG cells (PPiK(iH3.3K27M), PPW+iH3.3K27M) in ON, OFF and OFF-ON groups. **G.–H.** Representative images of mCherry expression by flow cytometry in live BT245 KO+iH3.3K27M **(G)** and PPiK(iH3.3K27M) **(H)** cells in ON, OFF and OFF-ON. **K.-L.** BT245 KO+iH3.3K27M **(K)** and PPiK(iH3.3K27M) **(L)** cells in ON, OFF, OFF-ON were fixed, permeabilized, and stained with an anti-H3K27M antibody conjugated with Alexa-488 in BT245 KO+iH3.3K27M or Alexa-405 in PPiK(iH3.3K27M) for intracellular flowcytometry and representative images are depicted. **I.-J.** Quantification of mCherry-positive cells corresponding to panel **(G-H)** and H3K27M-positive cells corresponding to panel **(K-L)**. The mCherry and H3K27M positivity thresholds were defined using the OFF population, with gates set such that 98% of OFF cells were classified as negative. The same thresholds were then applied to the corresponding ON and OFF–ON samples for quantification. Data represented as mean ± SEM. All data performed in three biological replicates and analyzed by one-way ANOVA. Significance is indicated as *p < 0.05, **p < 0.01, ***p < 0.001, and ns = not significant. **O.–P.** Western blot analysis for H3.3K27M and H4 as loading control in BT245 KO+iH3.3K27M and PPiK(iH3.3K27M) cells in ON, OFF, and OFF-ON.

To establish inducible and reversible human cells, we first evaluated the inducibility and reversibility of the iH3.3K27M system in HEK293T cells (Fig. S1A). Addition of dox to the cell culture media induced robust mCherry fluorescence, indicating oncohistone expression, whereas dox removal led to a gradual loss of signal over 2–5 days, consistent with suppression of H3K27M expression (Fig. S1A). Re-exposure to dox restored mCherry expression, inferring inducible and reversible control of H3.3K27M across ON, OFF, and OFF-ON (Fig. S1B). Next, we generated inducible and reversible human DMG cells by using previously CRISPR edited and described DIPG XIII KO and BT245 KO cells ^8^. Cells were transfected with the iH3.3K27M construct to generate human DMG cells hereafter termed BT245 KO+iH3.3K27M and DIPG XIII KO+iH3.3K27M. We also generated an additional murine DMG cell line by transfecting the previously described PPW cells with the iH3.3K27M construct ^22^, hereafter termed PPW+iH3.3K27M. Inducible and reversible expression of the oncohistone was inferred by microscopy using mCherry fluorescence under ON, OFF, and OFF-ON in human cells (Fig. 1E) and murine cells (Fig. 1F), respectively. Additional representative images for PPiK(iH3.3K27M) and PPW+iH3.3K27M cells, are shown in (Fig. S1 C, D). To confirm that changes in H3.3K27M expression did not affect luciferase expression from the LUC-p53 construct, we assessed luciferase protein levels under ON, OFF, and OFF-ON. Luciferase expression remained detectable across conditions in both PPiK and PPW+iH3.3K27M cells. We also generated murine neural progenitor cells NPCs (iH3.3K27M) from IUE-derived C57BL/6 model, in which we similarly demonstrated dox-dependent regulation of H3.3K27M expression (Fig. S1 F)

Furthermore, mCherry as a surrogate readout for oncohistone expression and H3.3K27M protein levels, were assessed by live- and intracellular flow cytometry in HEK293T+iH3.3K27M cells (Fig. S2A-D) as well as in human and murine iH3.3K27M cells (BT245 KO+iH3.3K27M and PPiK(iH3.3K27M)) (Fig. 1G–J) (Fig. 1K–N). Flow cytometry of live cells showed increased mCherry expression upon dox induction, loss of signal after dox removal, and restoration upon re-induction (Fig. 1I, J and Fig. S1B). Consistent with mCherry expression, intracellular flow cytometry using fluorophore-conjugated H3K27M specific antibodies confirmed dox-dependent H3.3K27M expression, with clear ON, OFF, and OFF-ON shifts relative to IgG isotype controls (Fig. 1M, N and Fig. S1D). Overlay plots comparing H3.3K27M staining with IgG isotype controls for each ON, OFF, and OFF-ON across inducible cell lines are shown in Fig. S2F-H. Western blot analysis further confirmed dox-dependent and reversible H3.3K27M expression in human and murine DMG cells (BT245 KO+iH3.3K27M and PPiK) (Fig. 1O–P).

Because approximately 20% of DMGs harbor H3.1K27M mutations, which are associated with reduced tumor aggressiveness and increased radiosensitivity ^23–25^, we also generated a dox-inducible H3.1K27M construct (Fig. S3A) to enable future comparative studies. Using the same approach, we established human (BT245 KO+iH3.1K27M) and murine (PPW+iH3.1K27M) cells and validated expression by mScarlet fluorescence microscopy (Fig. S3B-C) and H3.1K27M expression by western blot (Fig. S3D). In summary, we have provided valuable resources, which can be used for future studies to assess oncohistone dependent biological consequences in cells and tumors.

### H3.3K27M inhibition promotes differentiation and alters treatment response

To determine the biological consequences of H3.3K27M inhibition, we assessed neurosphere formation, and differentiation. The neurosphere formation (Fig. 2A), size (Fig. 2B) and numbers (Fig. 2C) were reduced upon dox removal, accompanied by morphological features of differentiation (Fig. 2A) like attachment to the plastic in cells where oncohistone expression was inhibited. Not surprisingly, elevated expression of differentiation markers, GFAP and EYA4 was observed in the OFF group (Fig. 2D). Notably, re-induction of H3.3K27M (OFF-ON) reduced GFAP and EYA4 expression to levels similar to the ON, indicating reversibility of the differentiation-associated phenotype (Fig. 2D). To determine whether H3.3K27M inhibition affects cell proliferation, PPW+iH3.3K27M cells were maintained under ON or OFF conditions. No significant difference in proliferation was observed after 5 days between ON and OFF cells (pvalue= 0.1066, not significant), although OFF cells showed a trend toward increased proliferation (Fig. 2E). This trend may in part reflect the removal of doxycycline from the culture medium rather than an effect of oncohistone inhibition. Then, to determine whether H3.3K27M inhibition affects treatment response, we compared proliferation in PPW+iH3.3K27M ON and OFF cells following radiation or ONC201(dordaviprone or Modeyso™) treatment. ONC201, a DDR2 inhibitor, which results in elevated TRAIL dependent cell death, has recently been FDA-approved for recurring DMGs and shown efficacy in H3K27M expressing cells, pre-clinical DMG models and also in the clinic ^5,26^. Interestingly radiotherapy when combined with oncohistone inhibition significantly reduced proliferation when compared to cells expressing H3K27M at 8 and 16 Gy (Fig. 2F). In contrast, ONC201 did not result in significant differences in proliferation between ON and OFF (Fig. S4A), suggesting that ONC201’s activity may be independent of H3.3K27M in DMG. We next evaluated whether combination of radiotherapy with ONC201 would synergize with oncohistone removal and potentially radio sensitize cells. Interestingly, a small trend towards radiosensitization of ONC201 was observed in the cells expressing the oncohistone when comparing its IC50 values, but not in cells where H3K27M expression was lost (Fig. S4 B, C). To summarize our findings, removal of the oncohistone drives neurosphere differentiation and reduces neurosphere formation and size. Additionally, cells lacking mutant histone 3 expression are more radio-sensitive, likely due to having differentiated, as previously reported by our group ^21^.

**Figure 2.**
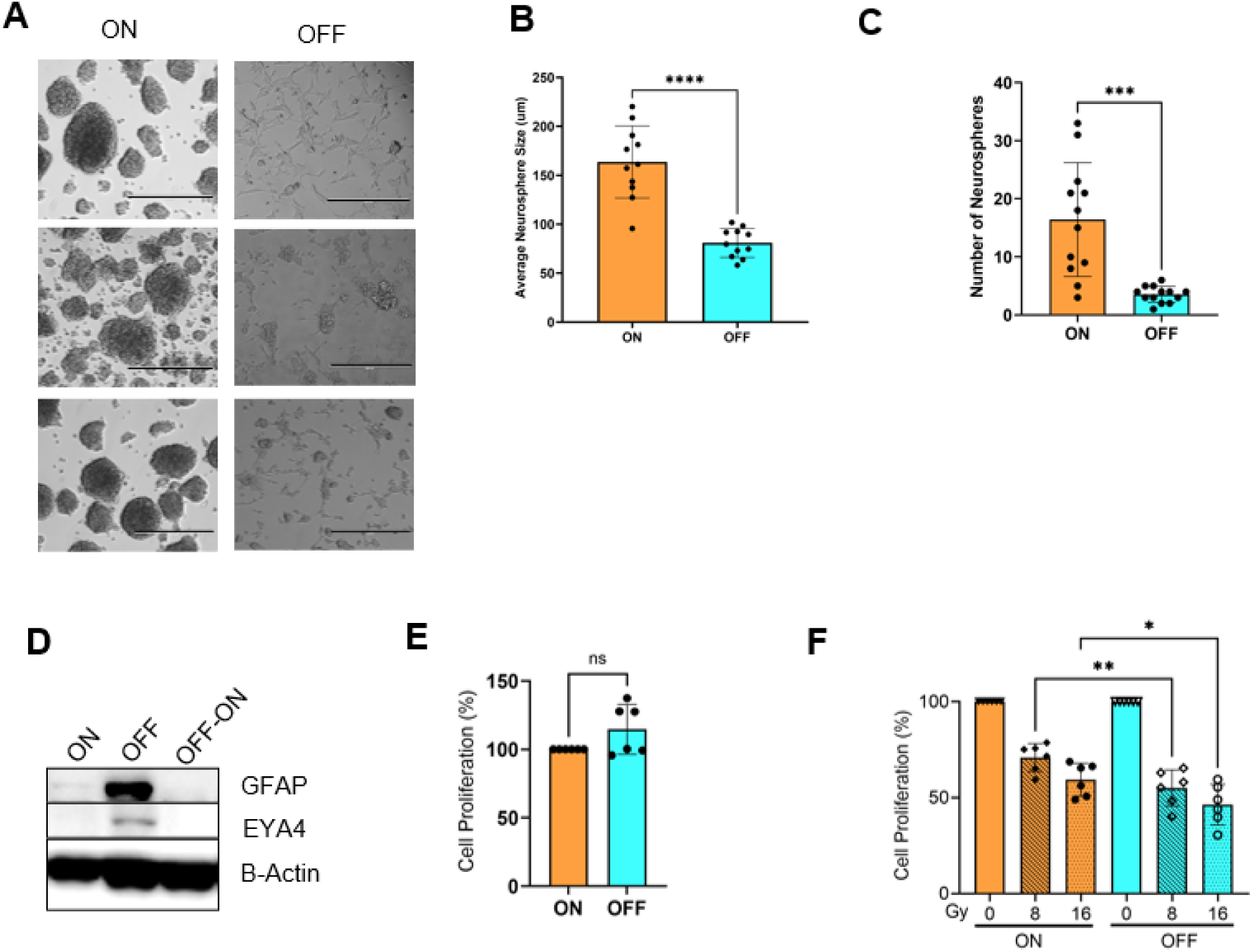
Biological consequences of H3.3K27M inhibition in DMG cells. **A.** Representative images of neurosphere formation on day 10 in PPiK(iH3.3K27M) cell in ON and OFF in three different biological experiments. **B.** Quantification of neurosphere size in ON and OFF at same timepoint described in **(A)**. Statistical analysis was performed using an unpaired two-tailed t-test, (****p < 0.0001). **C.** Quantification of neutrospheres’ number in ON and OFF at same timepoint described in **(A).** Statistical analysis was performed using an unpaired t test with Welch’s correction, (***p < 0.001). **D.** Western blot analysis of GFAP, EYA4 and β-Actin as loading control in PPiK cells in ON, OFF (day 17) and OFF-ON (day 5). **E.** Cell proliferation assay using TiterGlo of PPW+ iH3.3K27M cultured with and without dox 5 days after seeding. Statistical significance was assessed using a paired two-tailed t-test (ns =not significant). **F.** Cell proliferation assay using TiterGlo of PPW+ iH3.3K27M cultured with and without dox for 5 days after irradiation of 0, 8 or16 Gy. Statistical significance was assessed using one-way Anova (*p < 0.05, **p<0.01).

### H3.3K27M inhibition alters chromatin accessibility at the PDCD1 locus

To determine how H3.3K27M inhibition affects chromatin accessibility, we performed ATAC-seq in BT245 KO+iH3.3K27M cells under ON, OFF, and OFF-ON. ATAC-seq in BT245 KO+iH3.3K27M revealed changes in chromatin accessibility upon H3.3K27M inhibition (Fig. S5A). Annotation of accessible chromatin regions showed that the overall genomic distribution of ATAC-seq peaks was broadly similar across ON, OFF, OFF-ON (Fig. 3A). Intergenic regions represented the largest fraction of peaks in all groups, followed by intronic regions. Peaks at the promoter of transcriptional start sites (Promotor-TSS) showed only modest variation, while peaks at exons and TSS remained low and largely unchanged across all three groups (Fig. 3A). ATAC-seq analysis identified changes in chromatin accessibility between H3.3K27M ON and OFF cells, with partial restoration following H3.3K27M re-expression in the OFF-ON. Clustering of statistically and differentially accessible regions revealed clear difference between ON and OFF (Fig. 3B). Among the most prominent locus-specific changes, PDCD1 (PD-1) showed increased chromatin accessibility in the ON when compared to OFF (Fig. 3B). Genome browser tracks further confirmed these findings, demonstrating reduced ATAC-seq signal across the intronic regulatory region of PDCD1 following H3.3K27M inhibition, with partial restoration upon re-expression of H3.3K27M in the OFF-ON (Fig. 3C). To identify transcription factor motifs associated with chromatin regions that lose accessibility upon H3.3K27M inhibition, we performed motif enrichment analysis on ATAC-seq peaks that were decreased in OFF relative to ON. This analysis revealed enrichment of known motifs corresponding to OCT (octamer-binding transcription factor) family factors, PRDM15 (PR/SET domain 15), and Sp5 (Sp5 transcription factor) (Fig. 3D). A de novo motif corresponding to ZNF189 was also identified, though with lower confidence (P = 1 × 10) (Fig. 3D In summary, our chromatin accessibility study identified a limited number of differentially accessible regions, including increased accessibility at the PDCD1, CADM1 and CRB1 loci in the ON and motif analyses suggest that chromatin regions are enriched for specific transcription factor binding motifs upon oncohistone expression.

**Figure 3.**
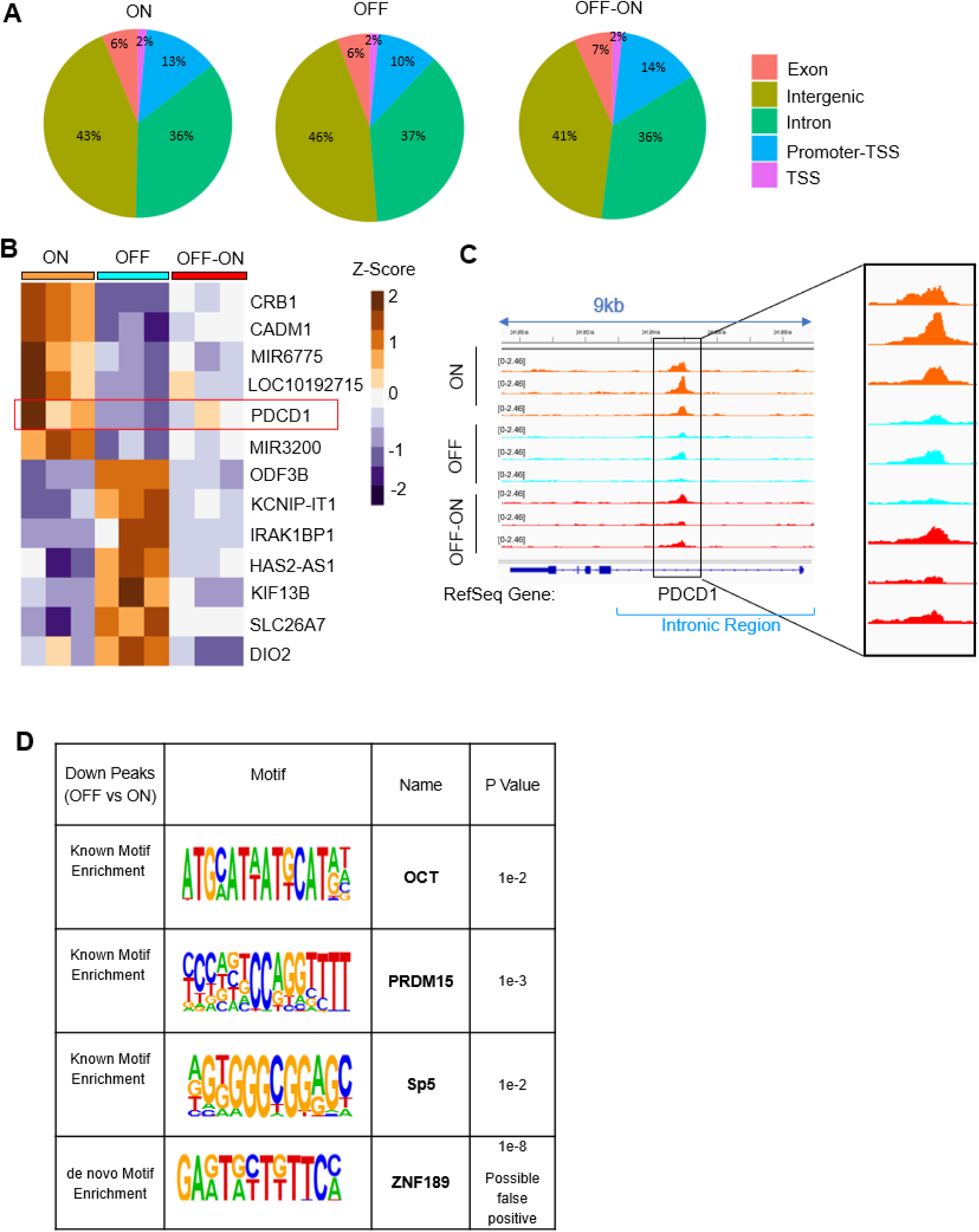
Reduction in chromatin accessibility upon H3.3K27M inhibition in DMG cells. **A.** Pie chart summarizing ATAC-seq peak distribution in BT245 KO+iH3.3K27M ON, OFF and OFF-ON. TSS refers to transcription start site. **B.** Heatmap showing significant differential ATAC-seq chromatin accessibility across ON, OFF, and OFF-ON in BT245 KO+iH3.3K27M cells. Differential analysis was performed using DESeq2, and regions were defined by |log FoldChange| ≥ 1 and FDR ≤ 0.05. Color scale represents normalized accessibility values (Z-score). **C.** Genome browser tracks at the PDCD1 locus in ON, OFF, and OFF-ON. Inset: peak magnification at the intronic region of PDCD1. **D.** Motif enrichment analysis of known and de novo ATAC-seq peaks using HOMER (Hypergeometric Optimization of Motif EnRichment) when comparing OFF to ON.

### Limited survival benefit of oncohistone inhibition in advanced tumors

To assess the inhibition of H3.3K27M expression in established tumors, we implanted BT245 KO+iH3.3K27M cells into the flank of immunocompromised mice and allowed them to grow until tumors reached approximately 50 cm³. At this point, mice were randomized to ON or OFF groups (Fig. 4A). Tumors in the ON group showed a consistent increase in tumor volume, whereas tumors in the OFF group, where oncohistone expression was turned off by removing dox from the chow, exhibited a decrease in tumor volume over time (Fig. 4B). The decreased tumor growth in the OFF group was maintained until collected at 6 weeks, when ‘ON’ tumors reached approximately 400 mm³ in volume (Fig. 4B). Analysis of tumor volume fold change showed a significant (p=0.0054) difference between ON and OFF groups (Fig. 4C). Western blot analysis of tumor tissues collected at the endpoint confirmed sustained H3.3K27M expression in ‘ON’ tumors and loss of H3.3K27M expression in OFF tumors (Fig. 4D). To assess whether inducibility and reversibility were retained after *in vivo* expansion, a tumor in the ON group was harvested and cells tested for H3K27M and mCherry expression. As expected, tumor-derived cells maintained their dox-dependent inducibility of mCherry expression assessed by microscopy, (Fig. S6A) and H3.3K27M expression by western blotting (Fig. S6 B, C). However, despite the described changes in H3.3K27M expression, no differences in H3 acetylation or methylation were observed in human and murine cell lines or tumor tissues (Fig. S6 D, E) indicating that epigenetic changes may be delayed when compared to oncohistone modulation.

**Figure 4:**
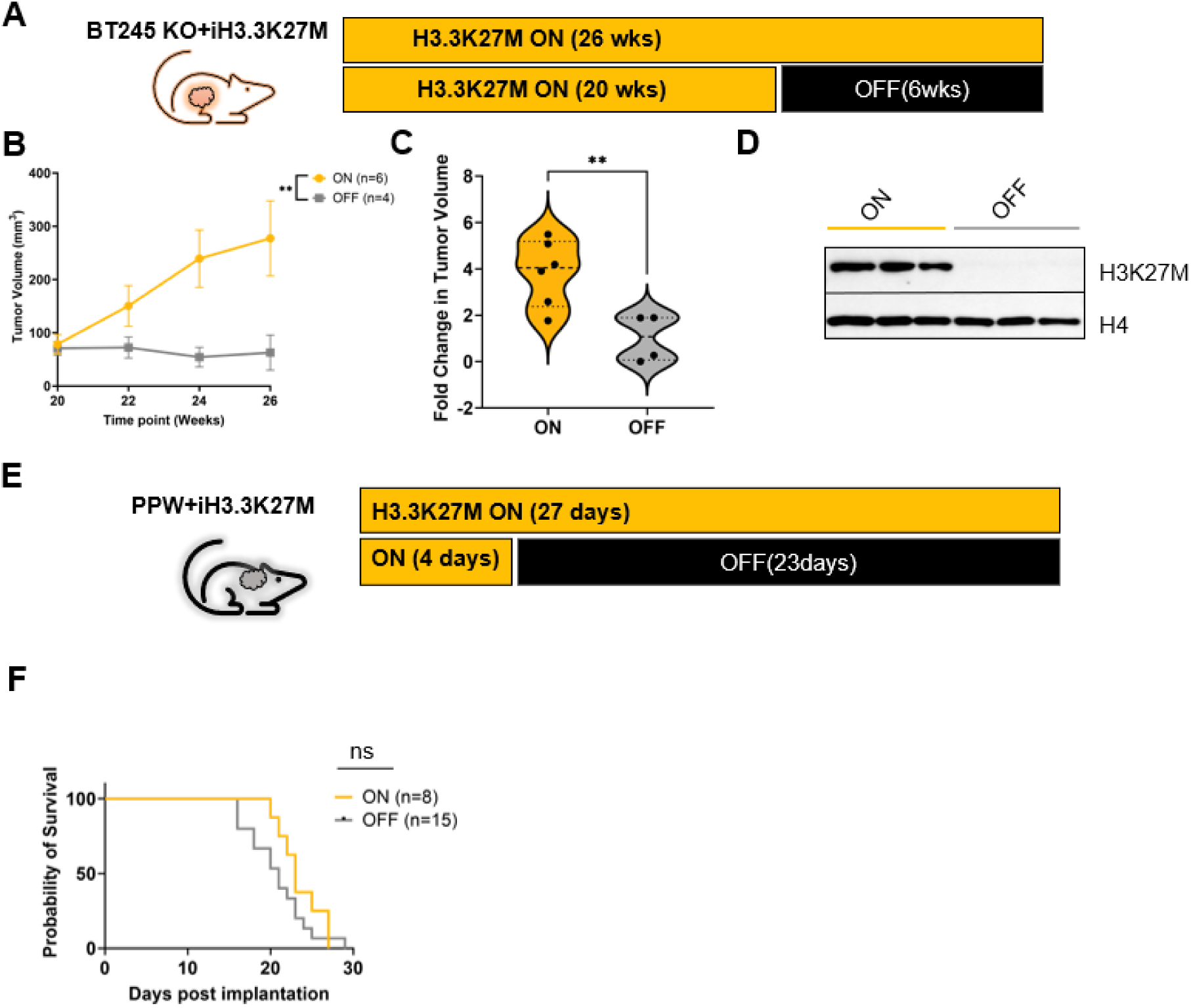
Efficient inhibition of H3K27M expression in tumors in flank and orthotopic xenograft H3.3K27M models. **A.** Experimental timeline: human BT245 KO+iH3.3K27M cells were implanted into the flank of NU/NU mice fed with dox for 20 weeks (ON) prior to removing dox (OFF) for 6 weeks. **B.** Longitudinal caliper measurements of BT245 KO+iH3.3K27M flank tumor volume over time (26 wks). **C.** Violin plot showing fold change in tumor volume at endpoint (26 wks) comparing ON and OFF groups in the BT245 KO+iH3.3K27M flank tumor normalized to the tumor size at the time of randomization. Each dot represents an individual tumor; horizontal lines indicate median values. Statistical analysis by unpaired two-tailed t test (**p < 0.01). **D.** Representative Western blot analysis of tumor lysates from BT245 KO+iH3.3K27M flank tumors in the ON (26 weeks) and OFF (6 weeks). Histone H4 serves as a loading control. **E.** Experimental timeline: murine PPW+iH3.3K27M were implanted intracranially into C57BL/6 mice. **F.** Kaplan–Meier survival analysis of PPW+iH3.3K27M tumor-bearing mice in the ON (n=8) or OFF (n = 15). Survival curves were compared using the log-rank (Mantel–Cox) test (ns=non-significant).

To assess the effect of H3.3K27M inhibition in an orthotopic model, PPW+iH3.3K27M cells were implanted intracranially into immunocompetent C57BL/6 mice and placed on a dox containing diet (Fig. 4E). After 4 days, mice were randomized into two groups and dox chow removed in the OFF group. However, surprisingly Kaplan–Meier survival analysis revealed no statistically significant difference between ON and OFF groups (Fig. 4F). These findings indicate that H3.3K27M inhibition alone may not suffice to impair tumor progression and expansion in the pons but may provide opportunities for combinatorial radiotherapeutic studies.

### H3K27M inhibition disrupts tumor–neuron crosstalk and reshapes neuronal signaling in DMG

To investigate the impact of H3.3K27M expression on tumor microenvironmental changes, and the consequences of genetic inhibition of H3.K27M, we utilized the IUE model as depicted in Figure 1B. Pregnant female mice were place on dox chow 24 hour prior to IUE procedure on embryos at E13.5-14.5. Mice remained on dox water until pups were born and until weaning when pups were placed on dox water. Mice in ON groups were on dox for 42 days total and in OFF group, 35 days prior to dox removal for 7 days. Ex vivo bioluminescence imaging (BLI) confirmed development of tumor and was used for locating precise position of the tumor prior to excision. Tumor was then fixed and sent for FLEX scRNA-seq (Fig. 5 A). Western blot analysis of intracranial tumor tissue confirmed inhibition of H3K27M expression in the OFF group, accompanied by loss of mCherry signal, indicating effective dox-dependent regulation of inducible H3.3K27M expression in the brain (Fig. S7A). Next, single-cell transcriptomic analysis was performed to characterize the tumor microenvironment under ON and OFF groups. Consistent with H3K27M inhibition, *H3f3a* mRNA expression showed a modest but statistically significant decrease in the OFF (Wilcoxon test, p = 0.0007) (Fig. S7B).

**Figure 5:**
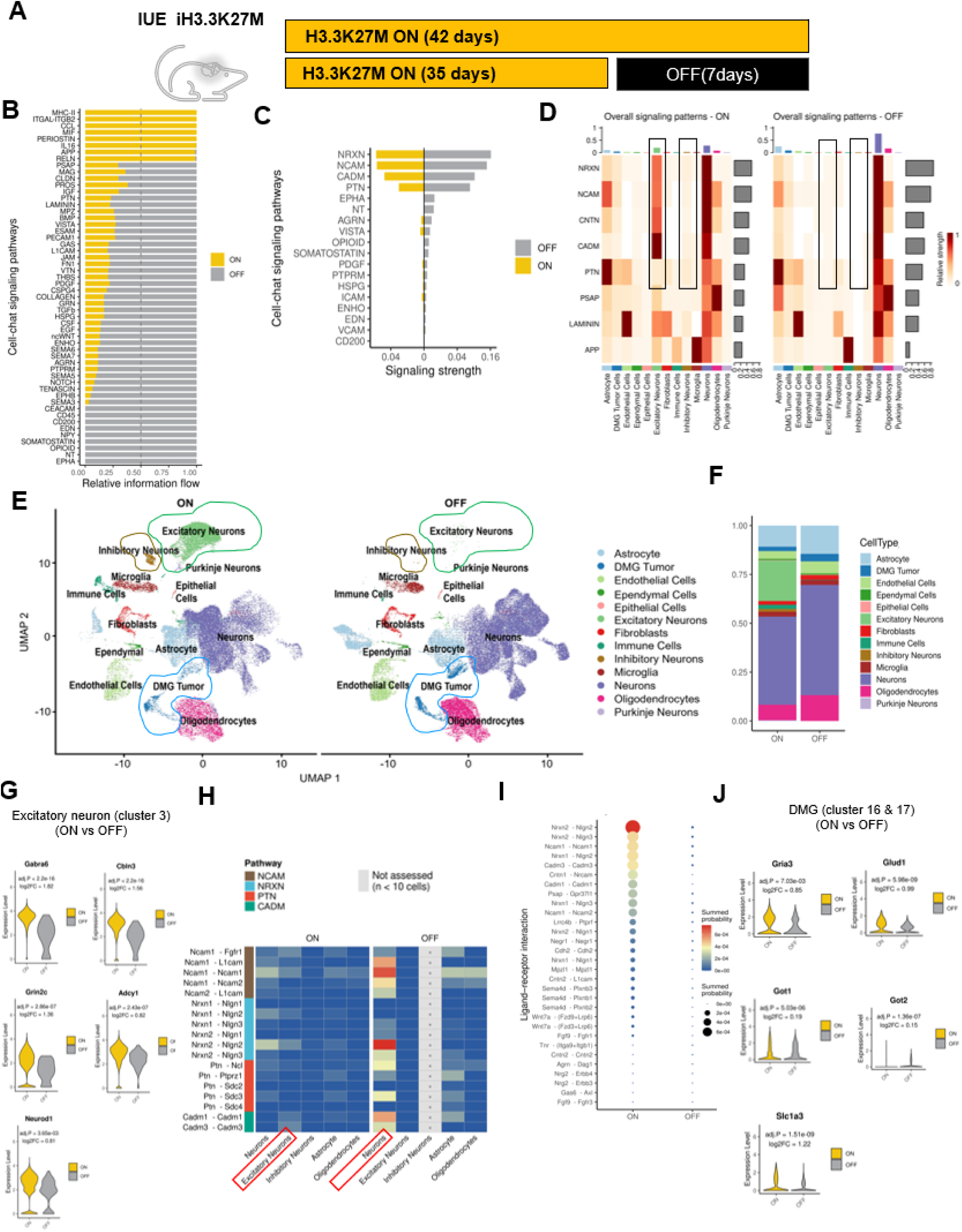
Identification of key signaling pathways (NRXN, NCAM, CADM, PTN) modulating signaling from DMG cells to excitatory neurons. **A.** Experimental design for inducible IUE H3.3K27M model: IUE procedure was performed as outlined in methods in CD1 mice and mice were fed dox containing chow starting at E14 until day 35. Mice were then randomized into two groups: ON (42 days total) and OFF (35 ON, 7 days OFF) prior to tissue fixation and Flex scRNA-sequencing of 1 female and 1 male mouse per group. **B.** Significant signaling pathways across ON and OFF groups in all cell clusters using CellChat. Pathways are ranked by relative information flow, representing overall signaling strength. **C.** Selected signaling pathway activity in ON and OFF groups in DMG cells. **D.** Heatmap of selected signaling pathways in ON and OFF in all cell clusters. **E.** UMAP visualization of all cell clusters identified by Flex single-cell RNA sequencing in ON and OFF groups. DMGs, Excitatory and inhibitory neurons are circled. An empty column was included for inhibitory neurons in OFF to indicate that no ligand–receptor interactions were detected due to the near absence of this population. **F.** Relative abundance of each cell population in ON and OFF groups. **G.** Violin plots of selected marker gene expression in excitatory neuron cluster (cluster 3) Adjusted p-values and log2 fold changes are indicated for each gene. **H.** Heatmaps of ligand–receptor interactions of outgoing signaling from DMG cells to recipient cell populations in ON and OFF groups. **I.** Dot plot showing excitatory neuron-to-DMG ligand–receptor interactions, with color and size representing interaction probability. All statistical significance were assessed using a paired Wilcoxon test. **J.** Violin plots of glutamate uptake and metabolism-associated genes in DMG cells (cluster 16,17) in ON and OFF. Adjusted p-values and log2 fold changes are indicated for each gene.

Cell–cell communication analysis using CellChat revealed both shared and H3K27M-dependent signaling pathways in all cells (Fig. 5B). In DMG cells, Neurexins (NRXN), Neural cell adhesion molecule (NCAM), Cell adhesion molecule (CADM), and Pleiotrophin (PTN), were found to be the dominant signaling pathways in both groups (ON and OFF) (Fig. 5C), however further analysis demonstrated that these pathways mediate interactions with excitatory and inhibitory neurons in the ON group, but much less when the oncohistone expression is lost, indicating a shift in signaling (Fig. 5D). PTN signaling was also prominent in astrocytes but remained relatively similar between ON and OFF groups (Fig. 5D). Consistent with these findings, excitatory (cluster 3) and inhibitory (cluster 20) neuronal cell populations were detected in ON but were markedly reduced in OFF when using UMAP analysis (Fig. 5 E) (Fig. S7C) while no major changes in the DMG cell clusters (cluster 16 and 17) were observed (Fig. S7D). Markers used for cluster annotation are provided in Supplementary Table 2. This dramatic reduction in excitatory and inhibitory neurons was further confirmed through cell-type composition analysis indicating a decreased abundance of excitatory neurons (cluster 3: 20.81% vs 0.12%) and inhibitory neurons (cluster 20: 1.43% vs 0.01%) (Fig. 5F). Consistent with these findings, genes associated with excitatory neuronal activity, including *GABRA6, CBLN3, GRIN2C, NEUROD1*, and *ADCY1* significantly increases in expression in ON compared to OFF (Fig. 5G). Some of the markers, including *GRM4, CBLN1* and *SYT2* in excitatory neurons (cluster 3) and *CNR1, TFAP2B* and *PVALB* in inhibitory neurons (cluster 20) showed a trend toward lower expression in OFF, although they did not reach statistical significance (Fig. S7E). To gain insight into the interaction changes between tumor cells (DMG) and neurons, we analyzed ligand–receptor communication between DMG cells (senders) and selected neuronal and glial cells (receivers) including neurons, excitatory and inhibitory neurons, astrocytes, and oligodendrocytes. Analysis of ligand–receptor interactions revealed a clear shift in DMG-derived signaling between ON and OFF groups. Oncohistone dependent (ON) DMG-specific signaling pathways NRXN, NCAM, PTN, and CADM were directed mainly toward neurons and excitatory neurons, with more limited signaling toward inhibitory neurons (Fig. 5H). Inhibitory neurons were nearly absent in OFF, and no ligand–receptor interactions were detected for this population (Fig. 5H). Contrary, upon oncohistone inhibition (OFF), this pattern shifted, with reduced engagement of excitatory and inhibitory neurons and a relative increase toward neurons (Fig. 5H). To further examine excitatory neuron–tumor interactions, we analyzed ligand– receptor communication between excitatory neurons (senders) and DMG cells (receivers). In the ON group, strong interactions were observed between these populations, whereas these interactions were reduced in the OFF group, indicating diminished neuron-to-tumor signaling (Fig. 5I). Notably, several interactions present in ON were lost in OFF, including low-probability but potentially targetable pathways such as FGF9–FGFR1 and SEMA4D–PLXNB1/2/3, suggesting neuron-to-tumor signaling mechanisms that may be therapeutically exploitable (Fig. 5I). These results suggest that DMG cells may hijack a neuronal niche within the tumor, potentially by disrupting the excitatory–inhibitory balance, increasing neuronal excitability and establishing synaptic connections that drive tumor proliferation.

Because DMG cells showed increased interactions with excitatory neurons through pathways associated with synaptic organization and neuron–tumor interactions, including NRXN, NCAM, CADM, and PTN-associated signaling, we next investigated whether these interactions were associated with glutamatergic neuronal excitation and we examined the expression of genes involved in glutamatergic signaling, glutamate uptake, and glutamine/glutamate metabolism in DMG cells. Glutamate released from activated presynaptic excitatory neurons into the synaptic cleft can also be sensed by surrounding cells within the tumor microenvironment ^27^. Violin plot analysis showed that genes involved in glutamate uptake and metabolism, including *GRIA3*, *GLUD1*, and *GOT1/2*, *SLC1A3*, were significantly higher in ON compared to OFF (Fig. 5J). Together, these findings suggest that enhanced DMG–neuron interactions in the ON are associated with excitation of neurons through increased glutamatergic signaling and synaptic organization.

### H3K27M inhibition alters intercellular communication and restores neuronal and immune cell interactions

Using single-cell RNA sequencing (scRNA-seq), we assessed transcriptional changes associated with H3.3K27M inhibition by analyzing the expression of stemness and differentiation markers in DMG cells (clusters 16 and 17) in ON and OFF. Analysis of stemness and differentiation markers showed a trend toward higher expression of stemness-associated genes in ON tumors and increased expression of differentiation markers in OFF tumors; however, these differences did not reach statistical significance, except for RNA Binding Fox-1 Homolog 3 (RBFOX3), which was significantly upregulated in OFF (Fig. S7F). To further investigate stemness-associated features, we examined the expression of aldehyde dehydrogenase (ALDH) isoforms in DMG cells (clusters 16 and 17), given our previous findings that ALDH human DIPG cells showed more proliferation, metabolic activity, and tumor-initiating capacity ^21^. ALDH1L1 was the only significantly altered isoform and was higher in ON tumors compared to OFF, while other ALDH isoforms showed variable, non-significant patterns (Fig. S7G). Together, these results suggest that short-term genetic inhibition of H3.3K27M expression does not broadly alter stemness/differentiation- or ALDH-associated expression, with the exception of RBFOX3 and ALDH1L1.

To investigate how H3.3K27M inhibition reshapes cellular communication, we performed CellChat analysis to compare global interaction networks between ON and OFF groups. Network analysis based on the top 20% of ligand–receptor pairs, revealed changes in interaction patterns and strength across cell populations between ON and OFF (Fig. 6A, B). Notably, several interactions were observed in the OFF group that were not detected in the ON, indicating the emergence of new cell–cell communication pathways following H3.3K27M inhibition. Analysis of all interactions in both conditions is presented in (Fig. S7H). When the oncohistone was inhibited, more signaling pathways were observed compared to ON (Fig. 6C). In OFF, increased signaling was evident within immune cell**s** (cluster 18), including CD200, CD45, and CEACAM pathways, and neurotrophin (NT), somatostatin, opioid, and ephrin (EPHA) pathways within neuronal clusters (0,1,4,7, 9, 10, 12, 13,15) (Fig. 6C). To further identify potential targets of increased immune signaling in the OFF group, we performed subclustering of immune cells based on selected cluster markers provided in Supplementary Table 3. (Fig. 6D) This analysis revealed distinct immune cell populations, including immunosuppressive tissue-resident macrophages, CD8^+^ T cells, regulatory T cells and B cells. Cell-type abundance analysis revealed a marked reduction in cell numbers in all immune subclusters in the OFF compared to ON (Fig. 6E). Several immune subclusters, including, CD8 T cells, regulatory T cells and B cells, were absent in OFF, indicating reduced immune diversity and heterogeneity following oncohistone inhibition (Fig. 6E). Based on cell-type abundance, the predominant immune populations remaining in OFF were immunosuppressive tissue-resident macrophages and dendritic cells, suggesting that these cell types may be primary recipients of CD200 and CD45-associated signaling.

**Figure 6.**
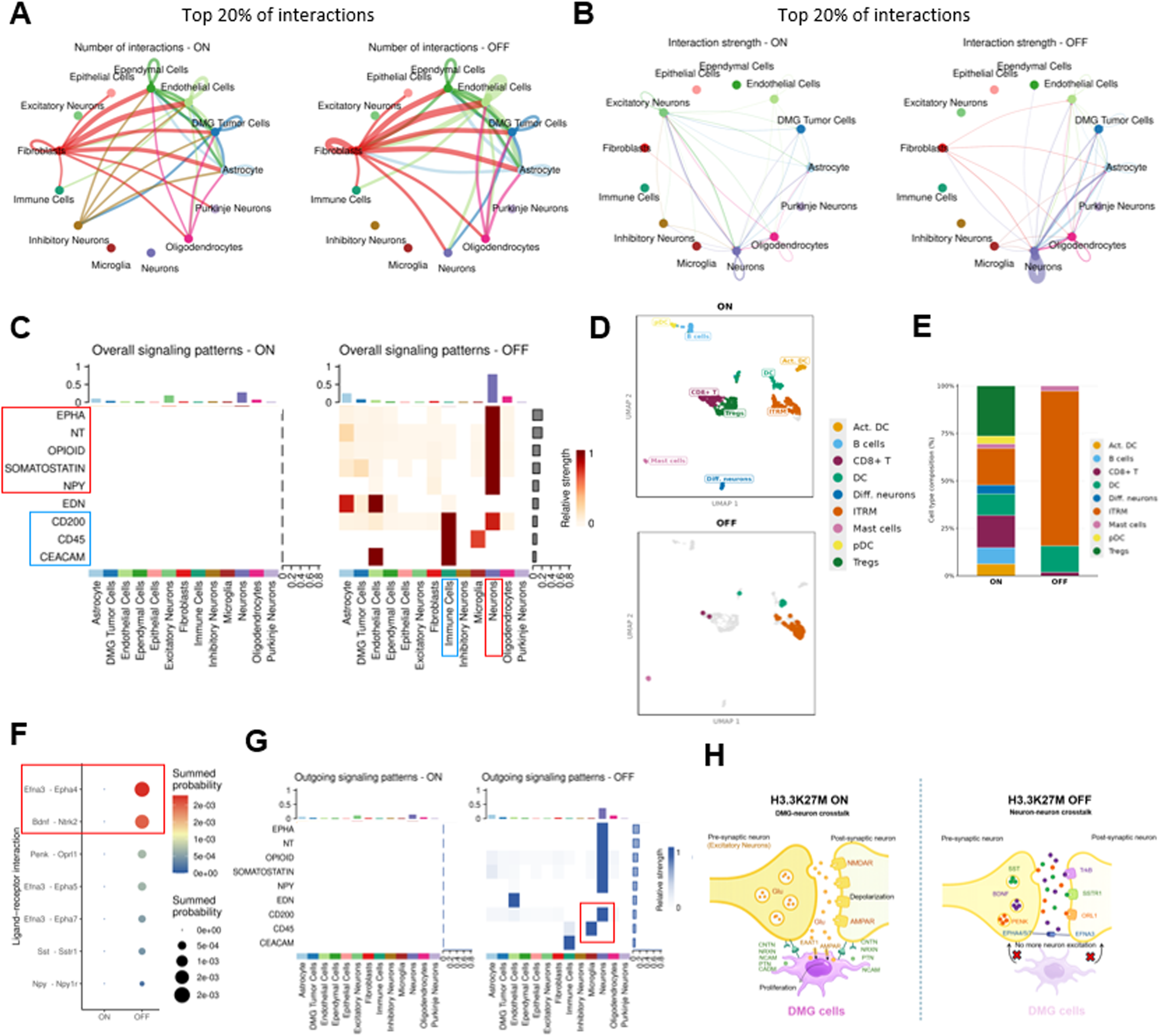
H3K27M inhibition alters communication networks in DMG. **A.** CellChat analysis of top 20% of ligand–receptor interactions in ON and OFF groups. **B.** Interaction strength of top 20% interactions in ON and OFF groups. **C.** Heatmap of signaling pathways specific to OFF group. **D.** UMAP visualization of immune cell subclustering in ON and OFF with annotated populations: activated dendritic cells (Act. DC), B cells, CD8 T cells, dendritic cells (DC), differentiated neurons, immunosuppressive tissue-resident macrophages (ITRM), mast cells, plasmacytoid dendritic cells (pDC), and regulatory T cells (Tregs). **E.** Relative abundance of immune cell subclusters in ON and OFF. **F.** Dot plot showing neuron-to-neuron ligand–receptor interactions identified in ON and OFF. Dot size represents summed communication probability. **G.** OFF-specific, outgoing signaling pathways for pathways represented in (**C**) in ON and OFF. **H. Schematic of DMG–neuron interactions in the H3.3K27M ON group and the shift to neuron-to-neuron communication with oncohistone inhibition**: DMG cells establish direct contact with neurons to establish synaptic connections through NRXN, NCAM, CADM and PTN pathways, leading to excitation of neuron. Activated pre-synaptic glutamatergic neurons synthesize and package glutamate into vesicles. Electrochemical stimulation of neurons results in release of glutamate in the synaptic cleft and activates post-synaptic receptors (NMDAR, AMPAR) leading to depolarization. Glutamate is taken up by DMG cells via EAAT1 and AMPAR. Glutamate uptake supports DMG cell proliferation. In contrast, H3.3K27M inhibition reduces DMG–neuron engagement and neuronal excitation, and more neuron-neuron communication is observed through EPHA, NT, Somatostatin, and NPY pathways with minimal involvement of DMG in these interactions. Abbreviations: AMPAR, α-amino-3-hydroxy-5methyl-4-isoxazole propionic acid Receptor; NMDAR, N-methyl-D-aspartate Receptor, EAAT1: Excitatory Amino Acid Transporter 1.

We next examined neuron to neuron interactions and identified ligand–receptor pairs associated with the enriched neuronal pathways observed in OFF in Figure 6C, including neurotrophin (NT), somatostatin, opioid, and ephrin (EPHA) signaling (Fig. 6F). Among these pathways, *EFNA3-EPHA4* (EPHA signaling) and *BDNF–NTRK2* (NT signaling) showed the strongest ligand–receptor interactions in the OFF. To further define the source and target of mentioned signaling in OFF, we analyzed outgoing (sender) signaling patterns across cell populations (Fig. 6G). Outgoing signaling analysis showed that CD200 signaling originated predominantly from neuronal populations and was directed toward immune cells (Fig. 6G). CD45-associated signaling was predominantly derived from microglia, and Endothelin (EDN) signaling was observed from endothelial cells toward astrocytes, indicating communication between vascular and glial compartments (Fig. 6G). Notably, with oncohistone inhibition, signaling was predominantly observed between neuronal populations, with minimal involvement of DMG tumor cells (Fig. 6G).

In summary, these results indicate that enhanced DMG–neuron interactions in H3.3K27M-associated groups are accompanied with increased neuronal excitation and glutamatergic signaling. After neuron excitation, glutamate will release into synaptic cleft and will be taken up by DMG cells via Excitatory Amino Acid Transporter 1 (EAAT1) coded by *SLC1A3* and α-amino-3-hydroxy-5methyl-4-isoxazole propionic acid Receptor (AMPAR) coded by *GRIA3* along with increased tumor proliferation. In contrast, upon oncohistone inhibition, DMG to neuron interactions are diminished, with reduced neuronal excitation and loss of tumor-associated signaling. With H3.3K27M inhibition, signaling is predominantly restricted and normalized to neuron–neuron interactions, with significantly reduced engagement of DMG cells (Fig. 6H).

## Discussion

Despite significant advances in understanding the role of H3K27M in DMG, limited therapeutic options exists and key questions remain regarding how its modulation influences tumor progression and whether oncohistone targeted approaches as future therapeutic options are warranted. Most current approaches to study H3.3K27M rely on comparative studies using isogenic cell lines and xenograft models thereof ^8,21,28,29^. While these studies have been instrumental in establishing the role of H3K27M in tumor initiation and epigenetic reprogramming, the consequences of oncohistone inhibition in clinically relevant tumor models have not yet been explored. However, such studies are critical for the development of future therapeutic strategies that directly target H3K27M with CRISPR/Cas9-based approaches, small molecule inhibitors or PROTACs as many therapeutic strategies targeting epigenetic modifiers or downstream signaling have been ineffective.

Therapeutic strategies directly targeting H3.3K27M are only beginning to emerge. Antisense oligonucleotides (ASOs) have shown promising preclinical activity, including induction of differentiation and reduced tumor growth ^9^. These studies highlight the therapeutic potential of targeting H3K27M in established tumors rather than preventing tumor initiation, as explored in implantation, GEMM, RCAS-TVA, and IUE models where oncohistone expression is omitted ^19,30,31^. However, these studies primarily evaluated early-stage disease in a single DMG model, leaving the effects of H3K27M inhibition in established tumors unclear. To address this gap, we developed inducible and reversible human, murine, and neural progenitor cell (NPC) H3.3K27M models, as well as inducible and reversible H3.1K27M models. These tools enable controlled H3K27M modulation during tumor progression, facilitate investigation of tumor-intrinsic and microenvironmental responses, and provide valuable resources for future comparative studies of H3.1- and H3.3-driven DMG biology in established tumors. Our findings further suggest that H3.3K27M extensively reshapes the tumor microenvironment, supporting direct oncohistone targeting as a potential therapeutic strategy for DMG. One limitation of this system is that complete oncohistone inhibition requires time following dox removal, which may affect therapeutic response kinetics in rapidly progressing tumors. Future studies will be required to optimize the timing and scheduling of H3K27M inhibition in established tumors.

The modest reduction in chromatin accessibility following genetic H3.3K27M inhibition is consistent with previous studies suggesting that H3K27M regulates selective epigenetic programs at specific loci rather than inducing widespread chromatin remodeling ^8,21^. Similarly, Harutyunyan et al. demonstrated focal alterations in H3K27 methylation patterns, while our previous Bru-seq study using isogenic H3K27M DMG cells identified broader transcriptional changes associated with H3K27M status ^8,21^. The relatively modest accessibility changes observed here may reflect the short duration of oncohistone inhibition (12 days OFF). Notably, several differentially accessible loci were linked to immune regulation and tumor– microenvironment interactions, including PDCD1 and HAS2-AS1, as well as cell adhesion and neuronal interaction pathways such as CADM1, KIF13B, and DIO2. In addition, reduced accessibility of motifs associated with OCT, PRDM15, Sp5, and a *de novo* motif associated with ZNF189 following genetic H3K27M inhibition is consistent with loss of stemness and chromatin regulation in DMG cells ^32–34^. Given the availability of PD-1/PD-L1-targeted therapies, these findings support future studies evaluating checkpoint blockade in combination with direct H3K27M inhibition, particularly in thalamic DMG models such as BT245 cells. Although immune checkpoint blockade has shown limited efficacy in DMG clinical and preclinical studies ^35,36^, it has not been evaluated in combination with direct genetic H3K27M inhibition, and future studies may determine whether similar epigenetic and transcriptional changes occur in pontine DMG models. Interestingly, despite efficient oncohistone inhibition, global H3K27 methylation and acetylation levels remained largely unchanged, suggesting that restoration of these epigenetic marks may require longer periods following H3K27M suppression in established tumors.

The limited survival benefit observed following genetic H3K27M inhibition in aggressive orthotopic tumors highlights the challenges of targeting the oncohistone in established DMG. Despite early H3.3K27M inhibition, rapid tumor growth and increasing tumor burden may limit therapeutic efficacy before complete oncohistone suppression is achieved. Because survival studies required long-term monitoring, we were unable to collect tumor tissue, and future studies will be needed to define the mechanisms underlying the limited survival response and to optimize the timing and duration of H3K27M inhibition. Importantly, the increased radiosensitivity observed following oncohistone inhibition suggests that radiotherapy may enhance the efficacy of H3K27M-targeted approaches by reducing tumor burden and providing a larger therapeutic window.

A major strength of our study lies in the use of scRNA-seq that enabled the investigation of genetic oncohistone inhibition within a developmentally and immunologically relevant brain microenvironment. Unlike orthotopic implantation models, which disrupt the native tumor microenvironment through direct injection into the brain, the IUE model enables tumor development within the intact developing brain and therefore may more closely recapitulate clinically relevant tumor–microenvironment interactions and changes ^18^. Because neuronal activity and neuron–glioma communication are key drivers of DMG progression ^37–39^, we specifically focused on tumor–neuron signaling and their response to oncohistone inhibition. We identified striking alterations in interactions between the tumor and excitatory neurons following genetic oncohistone inhibition. These findings are consistent with previous studies demonstrating that neuronal activity promotes glioma progression through both paracrine signaling and direct neuron-to-glioma synaptic communication ^39–41^. Our findings suggest that H3.3K27M promotes signaling programs associated with neuronal adhesion, synaptic organization, and tumor– excitatory neuron communication, including NRXN, NCAM, CADM, and PTN pathways ^42–44^. The pattern of NRXN, NCAM, and CADM signaling supports reciprocal cell–cell interactions between DMGs and excitatory neurons, whereas DMG-derived PTN ligand-receptor interactions support a distinct paracrine signaling mechanism toward excitatory neurons. These pathways have previously been implicated in neuron–glioma communication, neural niche remodeling, tumor invasion, and glioma progression ^41,42,44–46^.

In our study, Genetic oncohistone inhibition was associated with reduced immune cell abundance, which may reflect partial restoration of a more homeostatic brain microenvironment. Unlike peripheral tissues, the CNS normally contains few infiltrating lymphoid cells, with resident microglia representing the dominant immune population ^47^. Consistent with this, reduced T- and B-cell abundance following H3K27M inhibition may indicate decreased tumor-associated inflammatory signaling. Interestingly, Increased CD200-associated signaling further supports this interpretation, as CD200 helps maintain the immune-privileged state of the CNS by limiting excessive inflammation ^48^. After oncohistone inhibition, tissue-resident microglia and other immune cells actively communicate to maintain tissue homeostasis and respond to injury or stress caused by tumor cells. Therefore, despite reduced immune cell abundance, increased CD45-associated interactions suggest enhanced immune communication and homeostatic regulation within the brain microenvironment.

In parallel with the observed immune changes, genetic oncohistone inhibition was found to be associated with increased neuron–neuron interactions. These oncohistone inhibition-associated signaling, including neurotrophin (NT), somatostatin, opioid, ephrin (EPHA), and BDNF-associated signaling, are commonly linked to neuronal homeostasis, axon guidance, and mature neural network activity ^49–52^. Thus, these findings provide insight into how H3.3K27M-dependent signaling reshapes neuronal interactions within the DMG microenvironment to support tumor progression. Further studies will be required to validate these tumor–excitatory neuron interactions and define their mechanistic role in DMG progression. Importantly, several tumor– neuronal communication pathways identified in this study are therapeutically targetable. Among all enriched pathways in oncohistone active tumors, PTN signaling may be particularly attractive, as it can activate ALK and PI3K/mTOR pathways^53^, which can be targeted using small-molecule inhibitors such as ALK inhibitors (crizotinib or TAE684), or PI3K/mTOR pathway inhibitors (PI-103) ^54–58^. Our findings further suggest that DMG cells may rely on neuron-derived signaling pathways involving FGFR and SEMA4D-associated pathways, both of which have previously been implicated in glioma progression and therapy resistance ^59,60^. Corresponding inhibitors, including cordycepin and pepinemab, have already been explored ^61^. Future studies should evaluate whether combining H3K27M inhibition with these blood brain barrier-penetrant targeted therapies can further disrupt tumor-supportive communication within the brain microenvironment.

In summary, our inducible and reversible models provide insight into biological consequences of direct oncohistone targeting during tumor progression. Our H3K27M cell and mouse models provide resources to guide future therapeutic strategies, including CRISPR/Cas9-based and pharmacological approaches targeting H3K27M as we have demonstrated that genetic oncohistone inhibition not only affects intrinsic tumor cell signaling, but also extensively reshapes neuronal and immune interactions within the DMG microenvironment. Importantly, the differences observed between BT245, DIPGXIII, and murine DMG models suggest that the response to H3K27M inhibition may be cell- and model-specific. Having multiple inducible models allows us to identify which responses to H3K27M inhibition are shared across DMG models and which are cell- or model-specific, which will be important when considering H3K27M-targeted therapies. Ongoing studies will investigate the astrocyte–DMG, and myeloid– DMG crosstalk and their role in tumor progression and therapeutic response along with genetic oncohistone inhibition. While this study focused on H3.3K27M-driven DMG, future work may also evaluate whether similar tumor–microenvironment interactions and therapeutic vulnerabilities are present in H3.1K27M tumors.

## Credit authorship contribution statement

N.K: Investigation, Methodology, Formal analysis, Data curation, Writing; original draft and review & editing, M.I: Investigation, Formal analysis, Data curation, S.G: Investigation, Data curation, M.F: Investigation, Data curation, L.R: Investigation, Data curation, C.B: Investigation, Data curation, R.D: Investigation, Data curation, R.C: Investigation, Data curation, C.K: Writing; review & editing, Y.ZH: bioinformatics analysis, Writing; review & editing, S.G: Conceptualization, Methodology, Writing; original draft, Formal analysis, Writing; review & editing.

## Ethics

All animal procedures were approved by the Institutional Animal Care and Use Committee (IACUC) of University of Michigan and Unit for Laboratory Animal Medicine (ULAM) under approved protocol PRO00012162 to Stefanie Galban.

## Acknowledgements

Flow Cytometry work was performed at the University of Michigan Flow Cytometry Core (Ann Arbor, MI, USA). ATAC-seq assay and bioinformatic analysis of the ATAC-seq data were performed by Active Motif (Carlsbad, CA, USA). scRNA Seq and library prep was carried out in the Advanced Genomics Core at the University of Michigan (Ann Arbor, MI, USA). Research reported in this publication was supported by the University of Michigan Advanced Genomics Core, the UM Single Cell Spatial Analysis Program and the National Cancer Institutes of Health under Award Number P30CA046592 by the use of the following Cancer Center Shared Resource: Single Cell and Spatial Analysis Shared Resource.

We would like to thank Dr. Jabado’s lab (Pediatrics, McGill University) for providing isogenic pairs (human SU-DIPG XIII parental (H3.3K27M), CRISPR edited SU-DIPG XIII K27M-KO, and BT245 H3.3K27M parental and CRISPR edited BT245 K27M-KO) ^8^, and Dr. Timothy Pheonix (University of Cincinnati) for providing murine DMG cell lines (UC-BL6-B7(PPK) and UC-BL6-D3(PPW)) ^18^.

## Data availability

All analysis code and figure-generation scripts are publicly available on GitHub. The sequencing data generated in this study have been deposited in the Gene Expression Omnibus (GEO) database. The GEO accession number is pending and will be provided upon publication.

Supplementary Methods, Supplementary Figures, and Supplementary Table1,2,3 are provided.

