## Supplementary Figures for "Oncohistone inhibition reshapes tumor–microenvironment communication in Diffuse Midline Glioma (DMG)"

**Supplemental Figure Legends:**

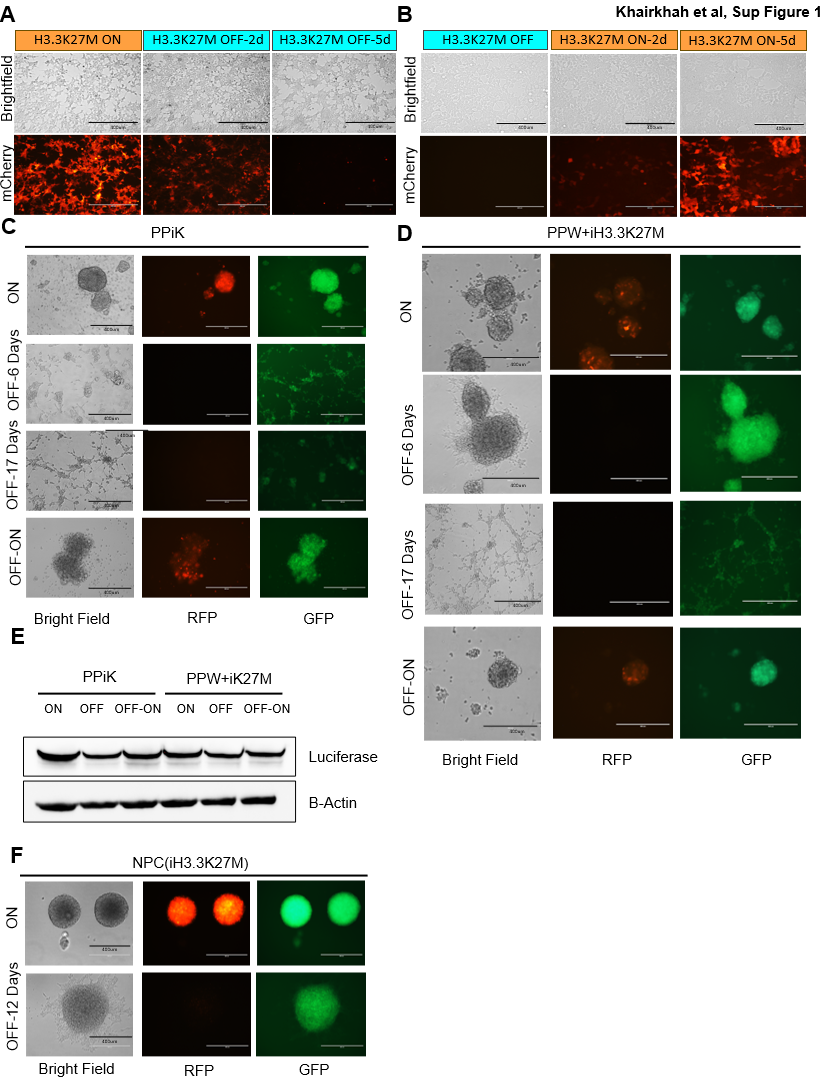

**Figure S1.** Validation of H3.3K27M inducibility and reversibility in cells and NPCs. A.-B. Representative phase contrast and mCherry fluorescence microscopy images in HEK293T+iH3.3K27M cells when cultured with dox (ON), when dox was removed for 2 and 5 days (OFF conditions) (A), and when dox was re-introduced to OFF condition for 2 and 5 days (B). C.-D. Representative microscopy imaging of mCherry and GFP (DNp53-IRES-GFP) expression in PPiK and PPW+iH3.1K27M cells treated with dox or when dox has been removed for 6 and 17 days (OFF) and when re-activated with dox (OFF-ON). E. Western blot analysis of luciferase expression in PPiK and PPW+iH3.3K27M cells in ON and OFF. F. Representative bright field and fluorescent images of murine neuronal precursor cells (NPCs) stably transfected with iH3.3K27M, generated from IUE C57BL/6 mouse. Scale bars for all microscopy images, 400 µm.

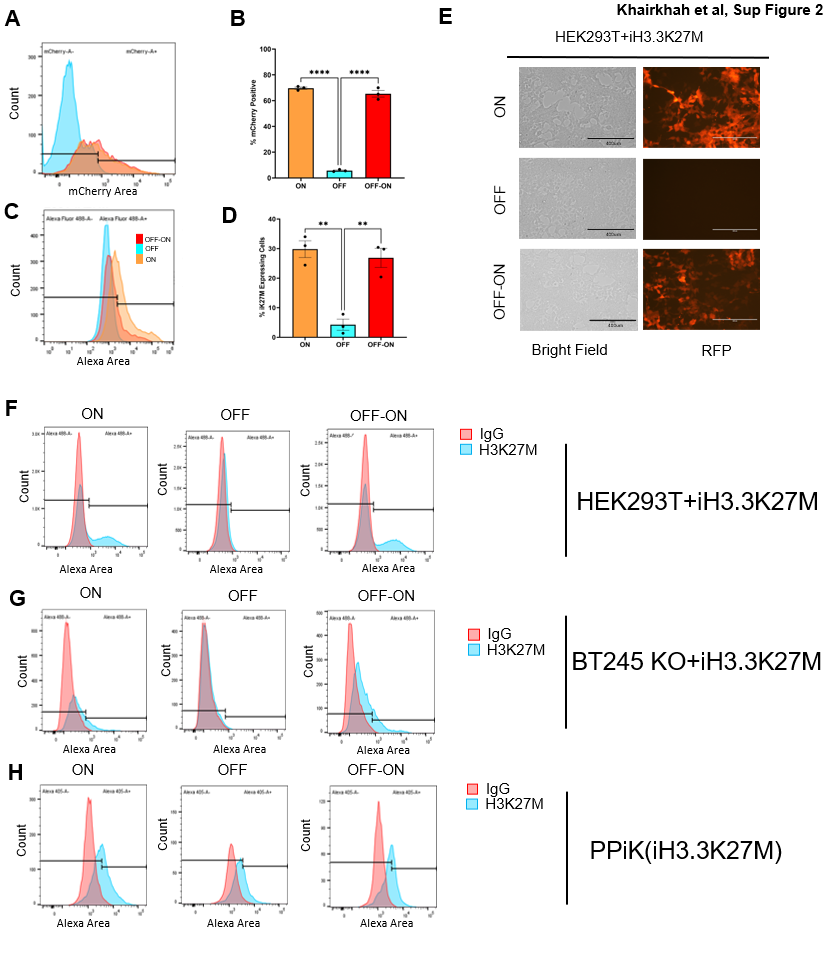

**Figure S2. Flow cytometry validation of inducible and reversible H3.3K27M expression in cells. A.-B.** Representative images of mCherry flow cytometry in live and **C.-D.** intracellular flowcytometry of H3.3K27M with Alexa-488 conjugated anti-H3K27M or IgG antibody in fixed and permeabilized HEK293T+iH3.3K27M cells and corresponding quantitative analysis. Data are shown as mean ± SEM in three biological replicates. Statistical analysis was performed with one-way Anova. ****p < 0.0001, **p < 0.01. **E.** Representative microscopy images of HEK293T+iH3.3K27M cells in ON, OFF, and OFF-ON corresponding to the flow cytometry results shown in panels **(A-D).** **F.-H.** Representative images of intracellular flow cytometry overlays of IgG isotype control (red) and H3K27M antibody (blue) staining in HEK293T+iH3.3K27M **(F)**, BT245 KO+iH3.3K27M **(G)** and PPiK(iH3.3K27M) **(H)** cells across ON, OFF, and OFF-ON conditions. Scale bars for all microscopy images, 400 µm.

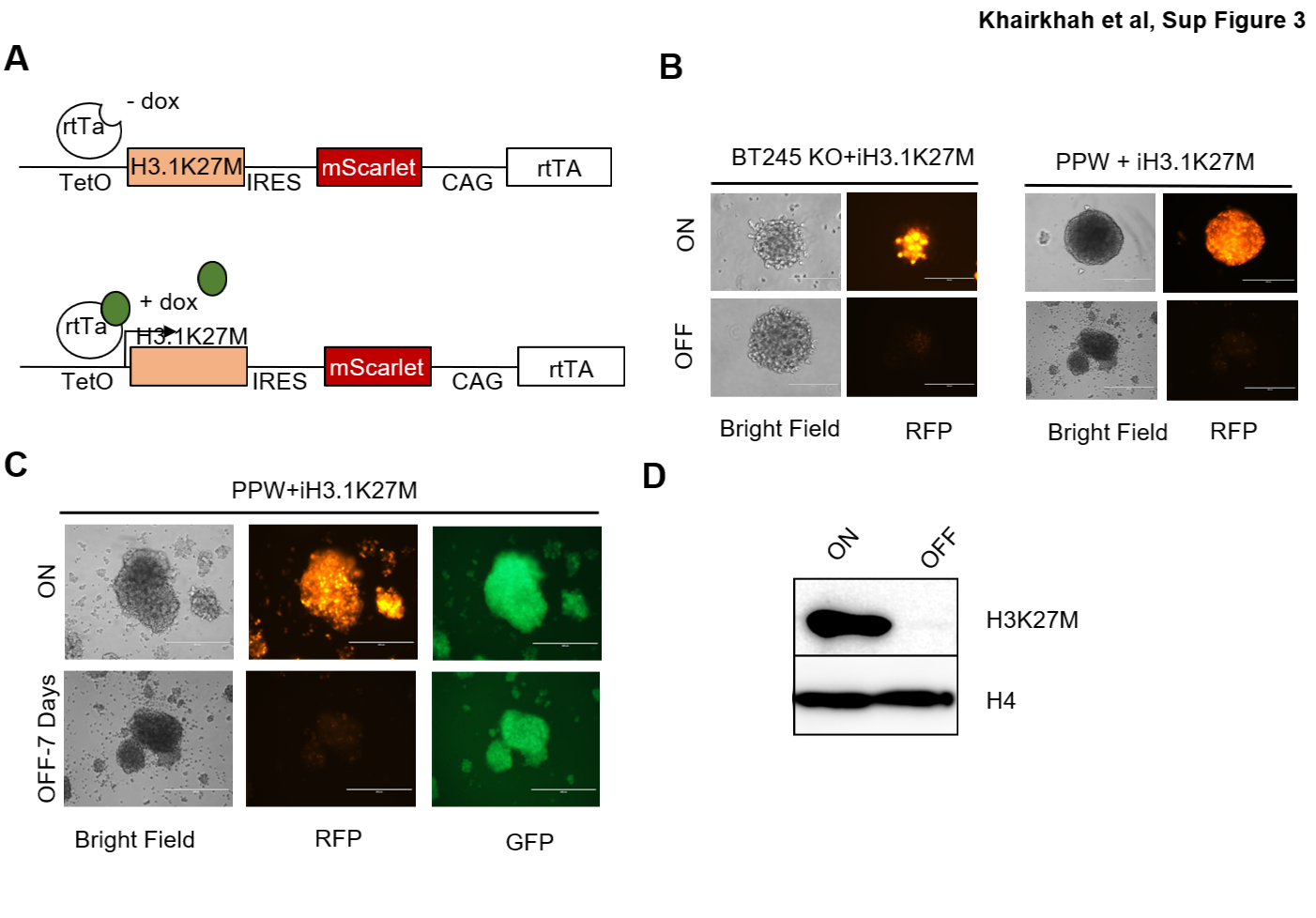

**Figure S3. Development of inducible and reversible H3.1K27M DMG cells.** **A.** Schematic of the Doxycycline-inducible H3.1K27M construct. In the presence of doxycycline (dox), rtTA can bind to the TetO element to induce expression of H3.1K27M and IRES dependent mScarlet reporter. **B.** Representative microscopy imaging of mScarlett expression in human (BT245 KO+iH3.1K27M), and murine DMG cells (PPW+iH3.1K27M) in ON (2ug/ml, dox) and OFF conditions. **C.** Representative microscopy imaging of mScarlet and GFP (DNp53-IRES-GFP) expression in PPW+iH3.1K27M cells treated with dox or when dox has been removed for 7 days (OFF). **D.** Western blot analysis of H3.1K27M and H4 expression as the loading control. Scale bars for all microscopy images, 400 µm.

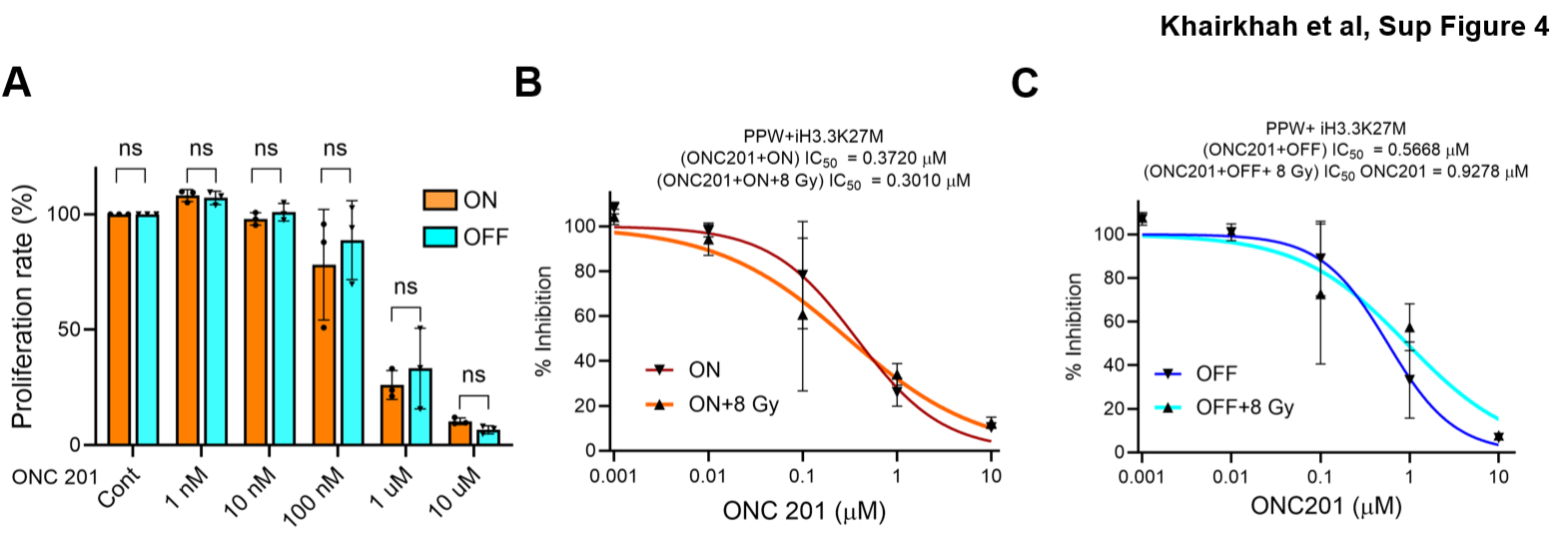

**Figure S4: Limited radiosensitization with Combined H3K27M Inhibition and Onc201. A.** Proliferation assay using TiterGlo in PPW+ iH3.3K27M cells cultured in ON or OFF conditions (+/- dox) and treated with increasing concentrations of Onc201 (1 nM–10 µM). Cell proliferation was measured relative to untreated controls. Statistical analysis was assessed using one-way Anova (ns, non-significant). **(B–C)** Dose–response curves showing percent inhibition and IC₅₀ values calculated from proliferation assays following treatment with Onc201 with or without 8 Gy irradiation in ON **(B)** and OFF **(C)** conditions.

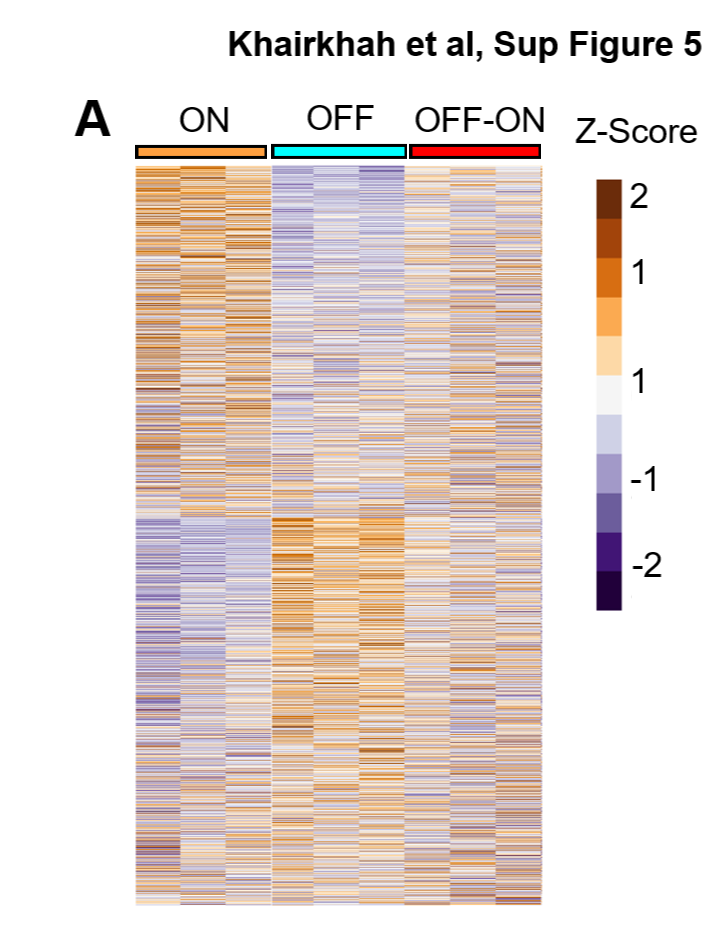

**Figure S5: Oncohistone inhibition results in global reduction of chromatin accessibility. A.** Heatmap depicting differential ATAC-seq chromatin accessibility of 1,000 regions across ON, OFF, and OFF-ON in BT245 KO+iH3.3K27M cells. Differential analysis was performed using DESeq2, and regions were defined by |log₂FoldChange| ≥ 1 and FDR ≤ 0.05. Color scale represents normalized accessibility values (Z-score).

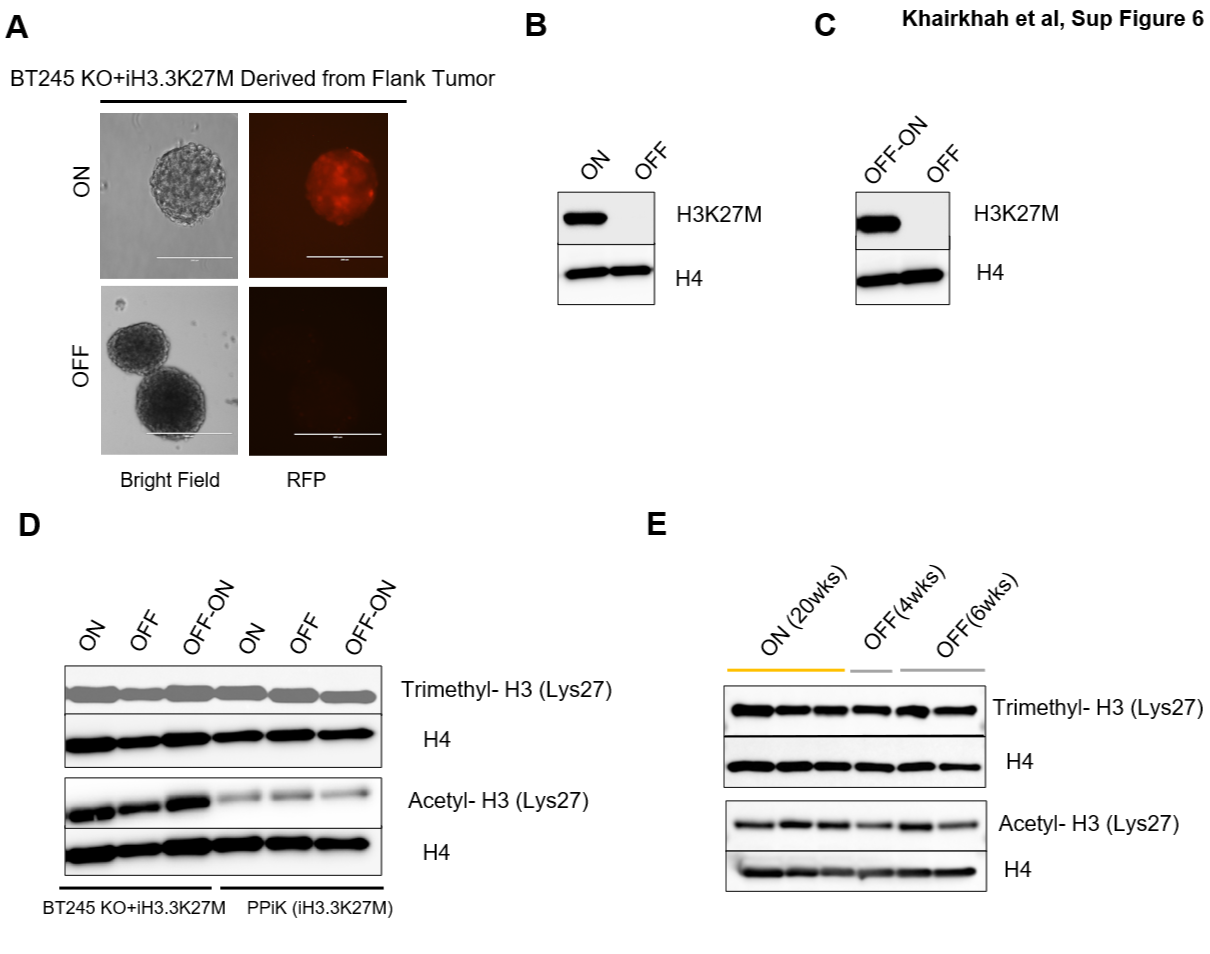

**Figure S6: Tumor–derived cells retain H3K27M inducibility and reversibility after long-term *in vivo* growth in flank xenograft model. A.** Representative bright-field and RFP fluorescence images of BT245 KO+iH3.3K27M cells derived from flank tumors in ON (20 weeks) or OFF (6 weeks) and subsequently cultured with (ON) and without (OFF) dox **B.** Representative western blotting of H3K27M expression and H4 as loading control in cells described in (**A**). **C.** Representative western blot of H3K27M expression and H4 as loading control in cells described in (**A**), but cells in OFF condition were re-exposed to dox (OFF-ON) and subsequently subjected to dox removal (OFF). **D.** Representative western blot of trimethylated H3K27 (H3K27me3), acetylated H3K27 (H3K27ac) and H4 as loading control in BT245 KO+iH3.3K27M and PPiK(iH3.3K27M) cells in ON, OFF, or OFF–ON conditions. **E.** Representative western blot H3K27me3, H3K27ac and H4 as loading control in tumor-derived BT245 KO+iH3.3K27M cells described in (**A**). Scale bars for all microscopy images, 400 µm.

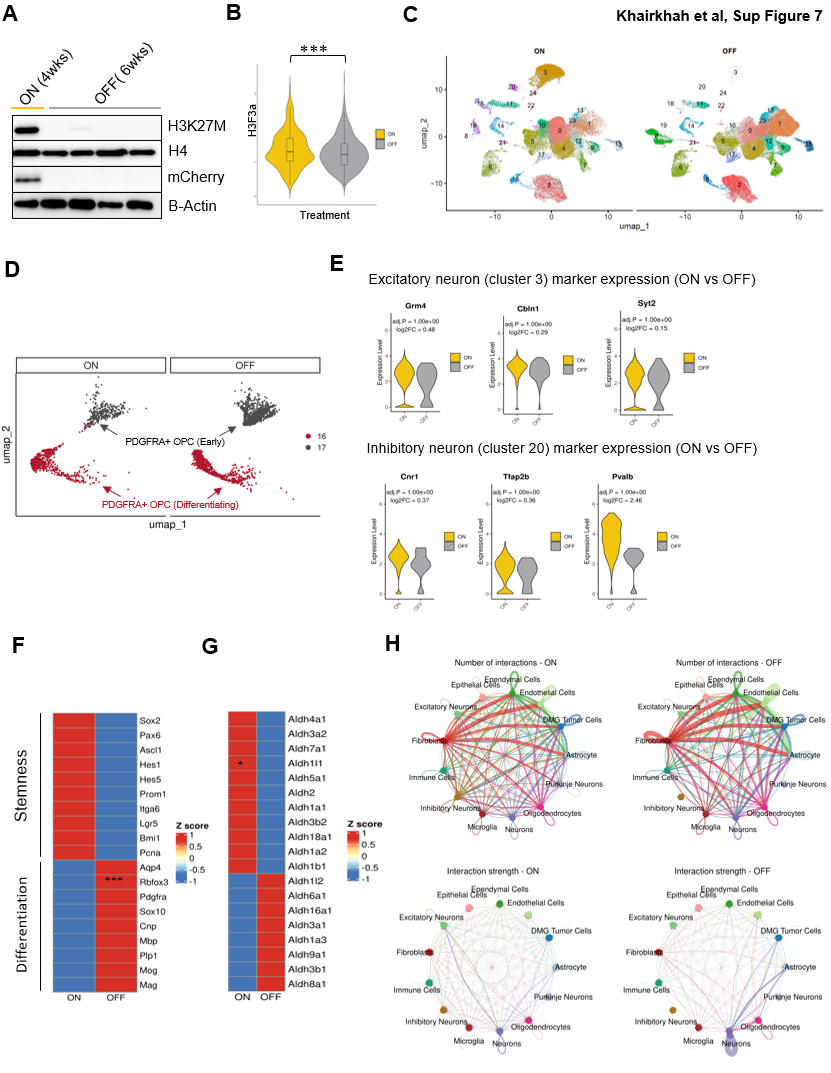

**Figure S7: H3K27M inhibition corrects neuronal and differentiation-associated marker expression. A.** Western blot analysis of H3K27M, mCherry and loading controls (H4 and β-actin) on tumor tissue from IUE DMG model (Fig. 5) in ON and OFF groups. **B.** Violin plot of H3f3a expression in ON and OFF from single-cell RNA-seq data (***p < 0.001). **C**. UMAP visualization of scRNA-seq clusters from H3.3K27M ON and OFF tumors. Cells are colored and numbered according to their identified clusters. Cluster-specific marker genes and corresponding cell-type annotations are provided in Supplementary Table 2. **D.** UMAP of DMG cells (PDGFRA⁺ OPC subclusters (clusters 16 and 17)) in ON and OFF. **E.** Violin plots of selected marker gene expression in excitatory neuron cluster (cluster 3) and inhibitory neuron cluster (cluster 20) in ON and OFF. Adjusted p-values and log2 fold changes are indicated for each gene. **F.** Heatmap of stemness/differentiation-associated marker expression in DMG cells in ON and OFF. Asterisk indicates statistically significant differential expression (***p < 0.001). **G.** Heatmap of ALDH isoform expression in DMG cells in ON and OFF. Asterisk indicates statistically significant differential expression (*p < 0.05). **H.** Number of cell-to-cell interactions and overall interaction strength in ON and OFF performed with CellChat. All statistical significance was assessed with Wilcoxon test.
