## Supplementary Methods for "Oncohistone inhibition reshapes tumor–microenvironment communication in Diffuse Midline Glioma (DMG)"

**Cell culture and culture conditions:**

The human DMG isogenic pair including human SU-DIPG XIII parental (H3.3K27M), CRISPR edited SU-DIPG XIII K27M-KO isogenic pair, and BT245 H3.3K27M parental and CRISPR edited BT245 K27M-KO, kindly provided by Dr. Jabado (Pediatrics, McGill University) (1), and the murine DMG cell line UC-BL6-B7(PPK) and UC-BL6-D3(PPW) cell lines, kindly provided by Dr. Timothy Pheonix (University of Cincinnati) (2) were used in this study. Human and murine cells were cultured as neurospheres as described previously (3). All cell lines were regularly checked for mycoplasma contamination using MycoAlert Mycoplasma Detection Kit (Lonza). Cell lines were cultured for a period of ∼ 2 months, after which they were replaced by new thaws.

**Generation of doxycycline-inducible H3.3K27M and H3.1 K27M vector:**

We designed a PiggyBac donor construct in which H3.3 K27M and mCherry , and H3.1 K27M and mScarlet are co-expressed via an internal ribosome entry site (IRES) under the control of a tetracycline-responsive element (TRE). The reverse tetracycline-controlled transactivator (rtTA) is encoded in the same plasmid and driven by a constitutive CAG promoter, enabling doxycycline (dox)-dependent activation H3.3 K27M-IRES-mCherry and H3.1 K27M-IRES-mScarlet cassettes. The pPB[TetOn]-TRE>H3.3K27M:IRES:mCherry-rev(CAG>tTS:T2A:rtTA) (Vector ID: VB240217-1015ebd) and pPB[TetOn]-TRE>{HIST1H3A H3.1 K27M}:IRES:Scarlet3-rev(CAG>tTS:T2A:rtTA) (Vector ID: VB250721-1078mgn) were synthesized by VectorBuilder (Chicago, IL, USA).

**Generation of inducible and reversible murine DMG cells using Intra uterine electroporation (IUE) model:**

The transposase helper plasmid pCAG-PBase, together with the donor constructs PB-CAG-PdgfraD824V-Ires-EGFP (PDGFRA D842V) and PB-CAG-DNp53-Ires-luciferase (dominant-negative TP53, referred to herein as TP53), were described previously and were used here without further modification (4). We also used PB TetON-TRE H3.3K27M-mCherry CAG rtTA to generate the inducible H3.3K27M IUE model. Dox (0.5 mg/mL) was added to the drinking water of pregnant CD1 mouse (The Jackson Laboratory) 24 hours prior to electroporation to initiate transgene activation in the developing embryos. Mice remained on dox-containing drinking water throughout the experiment until assigned to OFF, at which point dox was removed from the drinking water to inhibit H3.3K27M expression. IUE was performed at embryonic day E13.5-14.5 by injecting plasmid mixture into the lateral ventricles of embryos. Tumor formation was monitored by bioluminescence imaging (BLI). For *in vivo* bioluminescence imaging, mice were anesthetized using a 2.5% isofluorane/air mixture and injected with a single i.p. dose of 150 mg/kg Luciferin, Potassium Salt (Promega, WI). Consecutive images were acquired using an IVIS® imaging system (Perkin Elmer, Waltham MA). A tumor bearing mouse was harvested and processed to establish the PPiK tumors (P: PDGFRA, P: p53, iK: inducible H3.3K27M) cells.

**Generation of inducible and reversible human and murine DMG cells:**

The iH3.3K27M PiggyBac donor construct was first transfected into HEK293T and then DMG cells including SU-DIPG XIII K27M-KO (here DIPG XIII KO), BT245 K27M-KO (here BT245 KO) and PPW cells using Lipofectamine 3000 (ThermoFisher Scientific) according to the manufacturer’s protocol. Stable integration was achieved by co-transfection with the pCAG-PBase transposase plasmid in TSM complete medium supplemented with 2 μg/mL dox. mCherry-positive cells, indicating successful transfection and dox-induced transgene expression, were isolated by fluorescence-activated cell sorting (FACS) (BD FACS Discover S8 Cell Sorter at UMICH Flow Cytometry Core) at 72 h post-transfection and again one week later. Sorted cells were expanded and referred to as HEK293T+iH3.3K27M, DIPG XIII KO+iH3.3K27M, BT245 KO+iH3.3K27M, and PPW+iH3.3K27M.

**Live and Intracellular flow cytometry:**

We performed live and intracellular flow cytometry on three inducible cells, HEK293T+iH3.3K27M, BT245 KO+iH3.3K27M and PPiK(iH3.3K27M). HEK293T+iH3.3K27M and BT245 KO+iH3.3K27M cells were cultured under ON, OFF, and OFF–ON dox conditions. HEK293T+iH3.3K27M cells were maintained ON (+dox) for 7 days, OFF(-dox) for 5 days, or OFF–ON with 3 days OFF followed by 2 days ON. BT245 KO+iH3.3K27M cells were maintained ON for 12 days, OFF for 12 days, or OFF–ON with 7 days OFF followed by 5 days ON. The duration of H3K27M inhibition following doxycycline removal was optimized for each cell line based on cell line-specific responses and proliferation rates. Fixation and permeabilization were performed using the eBioscience Intracellular Fixation & Permeabilization Buffer Set (Thermo Fisher, Cat# 88-8824-00) one-step protocol for nuclear proteins. HEK293T+iH3.3K27M and BT245 KO+iH3.3K27M cells were stained with Histone H3 (K27M Mutant Specific) (D3B5T) Rabbit mAb, Alexa Fluor 488 conjugated, or Species-specific isotype control: Rabbit (DA1E) mAb IgG Isotype Control for 60 minutes at room temperature in the dark. PPiK (iH3.3K27M) cells were maintained under same conditions, with ON for 17 days, OFF for 17 days, and OFF–ON for 12 days OFF followed by 5 days ON. Due to spectral overlap between GFP and Alexa Fluor 488, intracellular staining for H3.3K27M in PPiK (iH3.3K27M) cells was performed using unconjugated primary antibody Histone H3 (K27M Mutant Specific) (D3B5T) Rabbit mAb or Rabbit (DA1E) mAb IgG Isotype Control, followed by incubation with goat anti-rabbit IgG (H+L) cross-adsorbed secondary antibody, Alexa Fluor 405. All antibody incubations were performed for 60 minutes at room temperature in the dark. Samples were acquired on ZE5 Cell Analyzer, 3 laser (405/488/561 nm) flow cytometer and analyzed using FlowJo v10.10. Manufacturer details for antibodies are listed in Supplemental Table 1.

**Western Blot:**

For protein extraction, two million cells were seeded 48h prior to collection and lysed with RIPA lysis buffer (Thermo Fisher) supplemented with protease inhibitors (Complete Protease Inhibitor Cocktail, Roche) and phosphatase inhibitors (PhosSTOP, Roche). For histone extraction, five million cells were seeded 48h prior to collection and followed by manufacturer protocol (Histone Extraction Kit Abcam). Protein concentration was determined using detergent compatible (DC) protein assay (Bio-Rad). Western blotting was performed as described previously (3). Ten micrograms of protein were separated on NuPAGE 12% Bis-Tris Acetate Protein gels (ThermoFischer Scientific) and transferred to a nitrocellulose membrane. Membranes were blocked with 5% skim milk, incubated overnight with primary antibodies followed by HRP-conjugated secondary antibodies. Details of all antibodies used in this study are provided in Supplementary Table 1. Enhanced chemiluminescence (ECL) substrate (Bio-Rad) and the Bio-Rad ChemiDoc MP imager were used according to the manufacturer’s recommendations.

**Neurosphere assay:**

PPiK(iH3.3K27M) cells were seeded as single-cell suspensions into 24-well at a density of 2×10^4^ and cultured in the ON and OFF conditions for 10 days. Fresh growth factors were added every three days. Dox was added to ON every three days, while an equivalent volume of water was added to OFF. We quantified neurosphere size and number using ImageJ software and averaged over the three fields of view, in three replicates, for three independent experiments.

**Cell proliferation assay:**

PPW+iH3.3K27M cells were seeded at 2,000 cells/well in 96-well plates and maintained under ON or OFF conditions (12 days OFF). Two hours after plating, cells were exposed to radiation at doses of 0, 8, or 16 Gy. Irradiation was carried out using Philips RT250 (Kimtron Medical) at indicated doses at the University of Michigan Comprehensive Cancer Center Experimental Irradiation Core. For drug treatment experiments, PPW+iH3.3K27M cells with same density were treated with increasing concentrations of ONC201(dordaviprone or Modeyso™) (Chimerix, WA, USA) (1nM,10nM, 100nM, 1µM and 10µM). For combination treatment, ONC201-treated cells were subsequently irradiated (8 Gy). In a separate experiment**,** cells were maintained under the same ON or OFF conditions without radiation. Cell viability was assessed 5 days later using the CellTiter-Glo assay (Promega), and luminescence was measured on an EnVision plate reader (PerkinElmer). All inhibitor assays were performed in triplicate. IC_50_ values were calculated using a nonlinear regression with variable slope by GraphPad Prism software (GraphPad Prism,)

**Bulk transposase-accessible chromatin with sequencing (ATAC-Seq):**

BT245 KO+iH3.3K27M cells were cultured under ON, OFF, and OFF–ON dox conditions same as before. Cells (250,000 per condition) were dissociated into single-cell suspensions, washed twice with 1X cold Phosphate Buffered Saline (PBS), and snap-frozen. Cryopreserved cells were sent to Active Motif (Carlsbad, CA) to perform the ATAC-seq. Frozen cell pellets were thawed in a 37 °C water bath, pelleted, washed with 1X cold PBS, and counted. Cells were lysed and tagmented using Tn5 transposase and reagents provided in the ATAC-Seq Kit (Active Motif), then PCR-amplified according to the manufacturer’s protocol. Libraries were cleaned using SPRI beads, quantified, and sequenced using paired-end 50-bp reads on an Illumina NextSeq 2000 platform. Bioinformatic analysis of ATAC-seq data was performed by Active Motif. Sequencing quality was assessed using FastQC (5), and adapter sequences were removed with Trim Galore. Trimmed reads were aligned to the human reference genome (hg38) using Bowtie2 (6). Low-quality reads, PCR duplicates, mitochondrial reads, multimapping reads, blacklisted regions, and improperly paired or excessively mismatched reads were filtered from further analysis using standard tools (Samtools (7), BEDTools (8), BAMTools (9), and Pysam (10)). Open chromatin regions were identified using MACS2 (11) in narrow peak mode with a false discovery rate (FDR) cutoff of 0.1. For visualization, aligned reads were converted to normalized bigWig files using deepTools (12) with counts-per-million (CPM) normalization. Differential chromatin accessibility was assessed using DESeq2 (13) based on consensus peak count matrices, with pairwise comparisons performed between OFF vs. ON, OFF-ON vs. OFF, and OFF-ON vs. ON conditions. Motif enrichment analysis of differentially accessible regions was performed using Hypergeometric Optimization of Motif EnRichment (HOMER) (14).

**Generation of inducible and reversible H3.3K27M *in vivo* models:**

All animal procedures were approved by the Institutional Animal Care and Use Committee (IACUC) of University of Michigan and Unit for Laboratory Animal Medicine (ULAM) under approved protocol PRO00012162 to Stefanie Galban.

8-12-week-old female NU/NU B/C nude mice (n=6) were obtained from Charles River Laboratories. Mice were bilaterally implanted with 5×10^6^ BT245 KO+iH3.3K27M cells per site, suspended in 50 µL of serum-free medium mixed 1:1 with Matrigel (Corning), subcutaneously into both flanks. All mice received dox-containing drinking water (0.5 mg/mL, in sterile water) continuously for 20 weeks to induce and maintain transgene expression. Tumor volume was measured weekly using digital calipers and calculated using the formula: volume = (length × width²) / 2. At week 20, when tumor volume reached 50 cm^3^, mice were randomized into two groups; (6 flanks per group): ON (+dox) and OFF (-dox). All mice were euthanized at week 26 when ON tumors reached approximately 400 mm³ in volume. At the endpoint, tumors were harvested, and tissue samples were snap frozen for further analysis.

Five- to eight-week-old female C57BL/6 (Charles River) mice (n=23) were used for intracranial implantation of murine PPW+iH3.3K27M cells. Mice were anesthetized and stereotactically implanted with PPW+iH3.3K27M cells into the brain as previously described (15). 300K cells were prepared as single-cell suspension in sterile phosphate-buffered saline and injected at a volume of 3µL per mouse using the coordination -0.8mm laterally, -1mm caudally and -4.5mm ventrally from the lambda. All mice received dox-containing drinking water (0.5 mg/mL, in sterile water) continuously 3days before implantation. After 4 days mice were randomized into two groups; ON (+dox) n=8 and OFF (-dox) n=15. Control mice remained on dox for the duration of the experiment. Mice were monitored for survival and were euthanized upon the onset of severe neurological symptoms.

**FLEX single cell RNA-sequencing (sc RNA-seq)**

In utero electroporation (IUE) was performed in CD1 embryos, using the same plasmid mixture and electroporation protocol described above. All moms received dox-containing drinking water (0.5 mg/mL, in sterile water). Pups were weaned at postnatal day 22 (P22) but remained with dox water and screened by BLI. Mice that remained BL-negative across two independent imaging sessions were euthanized. Once radiance reached approximately 10⁶ photons/sec/cm^2^/sr, mice were randomized into ON and OFF groups. After 7days, one male and one female in each group were euthanized and ex vivo BLI was performed immediately to localize tumors. Tumors and adjacent microenvironmental tissue were dissected within 5 minutes post-mortem and immediately snap frozen to send for FLEX single-cell RNA sequencing. University of Michigan Advanced Genomics Core processed snap frozen samples. Samples were thawed and immediately placed in fixation buffer from the Chromium Next GEM Single Cell Fixed RNA Sample Preparation Kit (10x Genomics), stored at 4°C overnight, and then Quenching Buffer was added according to the manufacturer’s protocol (10x Genomics, CG000553). Single-cell complementary DNA libraries were prepared and sequenced at the University of Michigan Advanced Genomics Core. Samples were run using 50-cycle, paired-end reads on the HiSeq 4000 (Illumina) to a depth of 100,000 reads. Raw FASTQ files were aligned to the mouse reference genome (mm10) and processed into feature-barcode matrices by the University of Michigan Advanced Genomics Core using the 10x Genomics Cell Ranger pipeline. Downstream single-cell analyses were performed using Seurat (v5.1.0) R package. Ambient RNA contamination was removed using DecontX (celda v1.20.0), and doublets were identified using scDblFinder (v1.18.0). Cells were filtered according to the following quality thresholds: genes detected per cell between 500 and 10,000 (nFeature), total UMI counts below 20,000 (nCount), and mitochondrial gene content below 5% (percent.mt). Data were log-normalized, highly variable genes were identified using the vst method, and scaled prior to principal component analysis (PCA). The first 12 principal components were used for downstream analyses based on elbow plot inspection.

**Sample Integration and Clustering and cell-type annotation**

To correct batch effects across individual samples while preserving biological variation, samples were integrated using Reciprocal PCA (RPCA) integration via Seurat's IntegrateLayers function. The four samples (NK13, NK14, NK15, NK16) were split by sample ID prior to integration. Clustering was performed using the Louvain algorithm (resolution = 0.4), and UMAP was used for visualization. Cell clusters were manually annotated based on the expression of canonical lineage marker genes. The marker genes used for each cluster, the cluster annotations, and the corresponding consolidated cell-type assignments are provided in Supplementary Table 2. To simplify downstream analyses and visualization, neuronal clusters with similar identities were grouped into a broader Neuron category, whereas excitatory and inhibitory neurons were retained as separate populations. Differentially expressed genes were identified using Bonferroni-corrected Wilcoxon rank-sum testing on the non-integrated RNA assay, with adjusted p-value < 0.05 considered significant.

**Differential gene expression analysis and Cell-Cell communication analysis**

Differential expressions between iH3.3K27M ON and OFF were assessed and results were filtered at an adjusted p-value threshold of < 0.05. Stemness and differentiation marker gene sets were visualized as heatmaps using average log2 fold-change values between ON and OFF conditions. Cell–cell communication analysis was performed using CellChat (v1.6.1) separately for ON and OFF conditions using the CellChatDB.mouse ligand–receptor database. Communication probabilities and signaling pathway activities were inferred and compared between conditions.

**Software and statistical analysis**

All analyses were performed in R v4.4.0. Statistical comparisons between conditions were performed using the Wilcoxon rank-sum test. Adjusted p-values were computed using Bonferroni correction unless otherwise specified. All analysis code and figure-generation scripts are publicly available on [GitHub](https://github.com/yzhao80/iH3.3K27M-DMG-scRNA-seq).
