## Supplementary Table 1 for "Oncohistone inhibition reshapes tumor–microenvironment communication in Diffuse Midline Glioma (DMG)"

| **Antibody Name** | **Company** | **Cat No** |  |
| --- | --- | --- | --- |
| Anti-Histone H3 (mutated K27M) antibody [EPR18340] - ChIP Grade | abcam | ab190631 | WB |
| Histone H4 antibody - ChIP Grade | abcam | ab10158 | WB |
| Histone H3 (K27M Mutant Specific) (D3B5T) Rabbit Monoclonal Antibody | Cell Signaling Technology | 74829 | FC |
| Rabbit (DA1E) Monoclonal Antibody IgG Isotype Control | Cell Signaling Technology | 3900 | FC |
| Goat anti-Rabbit IgG (H+L) Cross-Adsorbed Secondary Antibody, Alexa Fluor™ 405 | ThermoFisher Scientific | A-31556 | FC |
| Histone H3 (K27M Mutant Specific) (D3B5T) Rabbit Monoclonal Antibody (Alexa Fluor® 488 Conjugate) | Cell Signaling Technology | 85023 | FC |
| Rabbit (DA1E) Monoclonal Antibody IgG Isotype Control (Alexa Fluor® 488 Conjugate) | Cell Signaling Technology | 2975 | FC |
| Acetyl-Histone H3 (K27) (D5E4) XP® Rabbit mAb | Cell Signaling Technology | 8173S | WB |
| Anti-trimethyl-Histone H3 (Lys27) Antibody | Millipore Sigma | 07-449 | WB |
| GFAP (D1F4Q) XP® Rabbit mAb #12389 | Cell Signaling Technology | 12389S | WB |
| Anti-EYA4 antibody | abcam | ab251675 | WB |
| Secondary HRP-conjugated anti-rabbit IgG | Jackson ImmunoResearch | 111-035-003 | WB |
| Anti-Beta-Actin- HRP-conjugated | abcam | ab20272 | WB |
| Anti-Luciferase | Promega | G745A | WB |
