## Supplementary Table 2 for "Oncohistone inhibition reshapes tumor–microenvironment communication in Diffuse Midline Glioma (DMG)"

Supplemental Table 2 : Cell-type annotation of scRNA-seq clusters based on selected markers from the top 50 markers identified for each cluster.

| **Cluster #** | **Cell type** | **Top Selected Markers** | **Final name** |
| --- | --- | --- | --- |
| 0 | Mature Neurons | *Tubb3, Uchl1, Cplx1, Vamp1, Sncb, Nefl/Nefm/Nefh* | Neurons |
| 1 | Excitatory Cortical Neuron | *Camk2a, Nrgn, Stx1a, Kalrn, Ddn, Mef2c* | Neurons |
| 2 | Mature Oligodendrocytes | *Mog, Mal, Ugt8a, Fa2h, Gjb1 (Cx32), Gjc2 (Cx47), Tspan2, Hapln2* | Oligodendrocytes |
| 3 | Excitatory Neuron | *Gabra6, Cbln1 / Cbln3, Grm4, Grin2c, Neurod1, Syt2, Adcy1* | Excitatory Neuron |
| 4 | Cholinergic Neurons | *Ache, Nrsn2, Baiap3, Scg2, Caly, Cd200, Sema4g, Hap1* | Neurons |
| 5 | Malignant Glial | *Bcan, Cxcl14, Myorg, Ptprz1, Egfr* | Astrocyte |
| 6 | Vascular Endothelium | *Pecam1 (Cd31), Cldn5, Tek (Tie2), Ptprb (Ve-Ptp), Esam, Eng, Emcn, Adgrl4.* | Endothelial Cells |
| 7 | Striatal Medium Spiny Neuron | *Gpr88, Rgs9, Pde10a, Ppp1r1b (Darpp-32), Adcy5, Gng7, Penk, Rasd2* | Neurons |
| 8 | Reactive/Perivascular Astrocytes | *Aqp4, Mlc1, Slc38a3, Ntsr2, Fgfr3, Vcam1* | Astrocyte |
| 9 | Mature Excitatory Neurons | *Slc17a6, Ntng1, Lrrtm1, Slitrk6, Adarb1* | Neurons |
| 10 | Mature Neurons | *Snap25, Tuba1a/Tubb4a/Tuba1a, Eef1a2, L1cam, Rnf157* | Neurons |
| 11 | Microglia | *Tmem119,Trem2, P2ry13, Pld4, Siglech, Csf1r, Inpp5d, C1qa/C1qb* | Microglia |
| 12 | Inhibitory Interneurons | *Gad1, Gad2,Slc32a1* | Neurons |
| 13 | Mature Excitatory Neurons | *Neurod2, Neurod6, Grin2a* | Neurons |
| 14 | Fibroblasts | *Col1a1, Col1a2, Col3a1, Pcolce, Fmod, Ogn, Fbln1, Itih2, Sned1, Aebp1* | Fibroblasts |
| 15 | Neurons | *Icam5, Chrm1, Slit1, Synpr, Lrrtm4, Epha7, Sema5a, Pcdh8/Pcdh20* | Neurons |
| 16 | PDGFRA+ OPC (Differentiating) | *Pdgfra, Cspg4 (Ng2), Olig2, Sox10, Mki67, Trp53* | Dmg Tumor Cells |
| 17 | PDGFRA+ OPC (Early) | *Pdgfra, Cspg4 (Ng2), Olig2, Trp53* | Dmg Tumor Cells |
| 18 | Antigen-Presenting Myeloid Cells (Microglia/Macrophages) | *Ptprc (CD45), Itgb2, Il2rg, Antigen Presentation (MHC II): Cd74, H2-Aa, Tap1* | Immune Cells |
| 19 | Astrocyte | *Hopx, Mlc1, Kcnj16, A2m, Pltp, Cyp2j9, Lcat, Pcp2, Car8* | Astrocyte |
| 20 | Inhibitory Neurons | *Cnr1, Tfap2b, Pvalb* | Inhibitory Neurons |
| 21 | Ependymal | *Foxj1, Cd24a* | Ependymal Cells |
| 22 | Choroid Plexus Epithelial Cells | *Mfrp, Folr1, Kcnj13, Clic6, Tmem72, Slc4a5, Cldn2, Aqp1* | Epithelial Cells |
| 23 | Perivascular Fibroblasts | *Dcn, Col6a1/Col6a2, Col8a2, Fbln1, Emilin1, Efemp1, Olfml2a, Lox, Mgp* | Fibroblasts |
| 24 | Cerebellar Purkinje Neurons | *Pcp2 (L7)* | Purkinje Neurons |

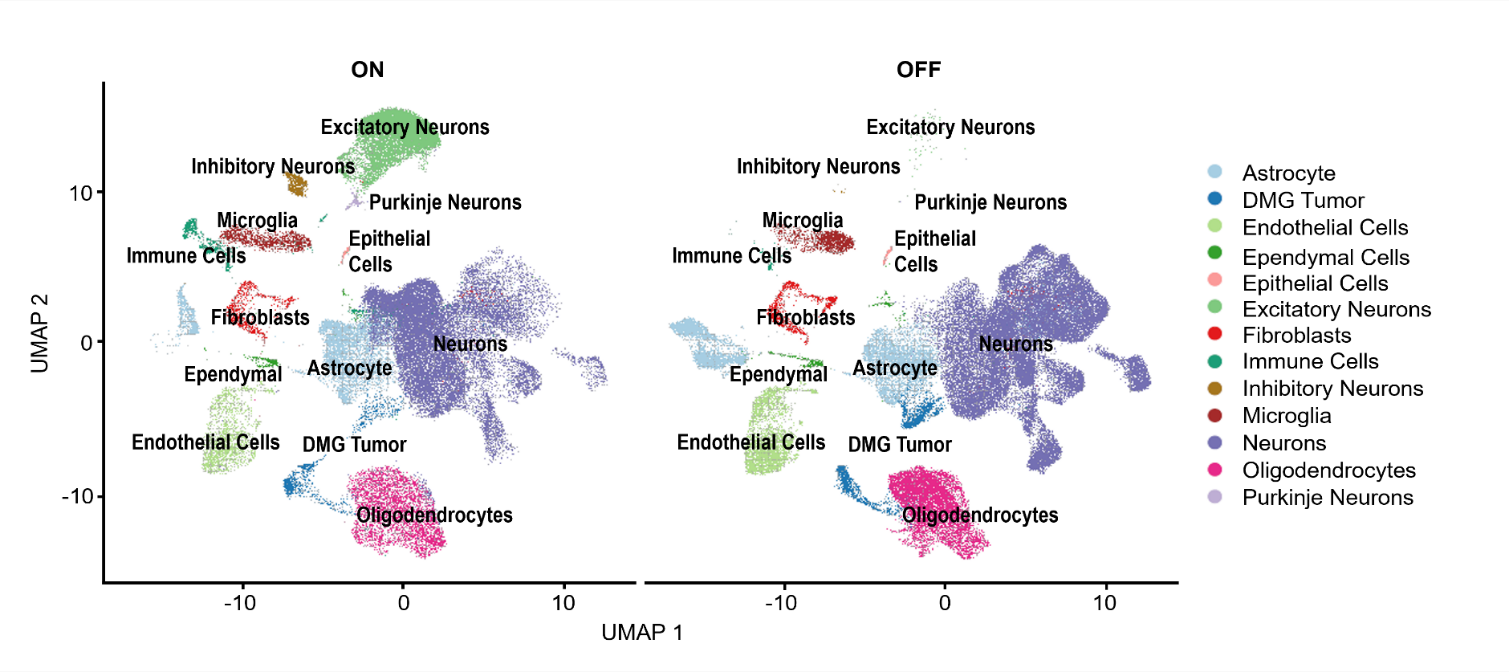
