## Supplementary Table 3 for "Oncohistone inhibition reshapes tumor–microenvironment communication in Diffuse Midline Glioma (DMG)"

Supplemental Table 3: Immune cell (Cluster 18) subcluster annotation based on selected markers from the top 50 markers identified for each subcluster.

| **Cluster #** | **Cell type** | **Top Selected Markers** | **Final name** |
| --- | --- | --- | --- |
| 0 | Neurons | *Tubb3, Nefl, Uchl1, Cplx1, Map2, Syp, Syn1, Eno2* | Contaminated Neurons (Will be removed) |
| 1 | Immunosuppressive tissue-resident macrophages | *Mrc1, Cd163, Msr1, Stab1, Mertk, Lyve1, Pf4, F13a1* | Immunosuppressive tissue-resident macrophages |
| 2 | T-Reg | *Foxp3, Il2ra, Ctla4, Icos, Tnfrsf4, Tnfrsf18, Ccr8, Ikzf2, Pdcd1* | T-Reg |
| 3 | CD8+ T Cell | *Cd8a, Cd8b1, Klrd1, Klrc1, Klrk1, Tbx21, Ccl5, Xcl1* | CD8+ T Cell |
| 4 | Dendritic Cell | *Xcr1, Clec9a, Wdfy4, Itgax, Tlr3, H2-Ab1, H2-Eb1, Ciita* | Dendritic Cell |
| 5 | B Cell | *Cd79a, Cd79b, Ms4a1, Cd19, Cd22, Pax5, Ebf1, Cd72* | B Cell |
| 6 | Activated Dendritic Cell | *Ccr7, Fscn1, Relb, Ccl22, Il12b, Il4i1, Ly75, Batf3* | Activated Dendritic Cell |
| 7 | Differentiated Neurons | *Gabra6, Cbln3, Grin2c, Grm4, Neurod1, Cbln1, Adcy1, Reln* | Differentiated Neurons |
| 8 | Plasmacytoid Dendritic Cells | *Siglech, Bst2, Gapt, Spib, Irf8, Ccr9, Scimp, Lair1* | Plasmacytoid Dendritic Cells |
| 9 | Mast Cells | *Tpsab1, Tpsb2, Cpa3, Kit, Ms4a2, Fcer1a, Mcpt4, Cma1* | Mast Cells |

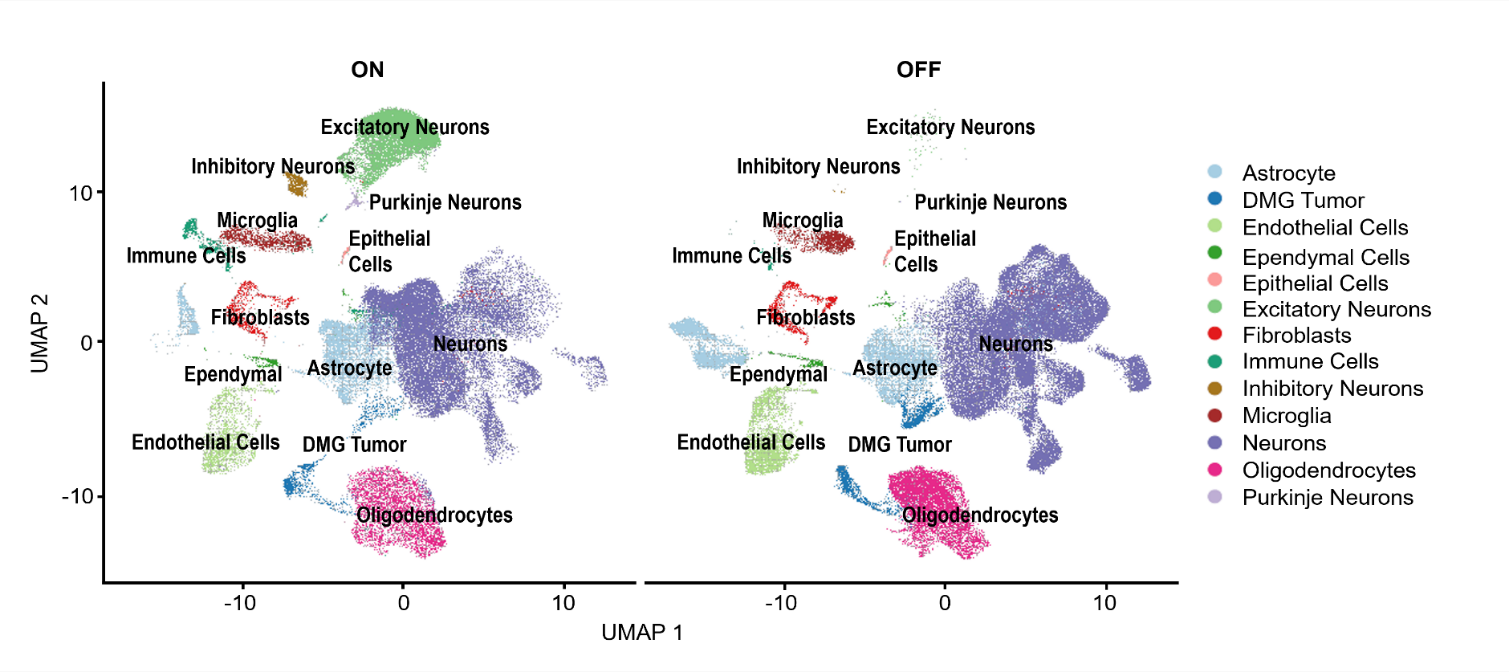
